# Bacterial origin of thymidylate and folate metabolism in Asgard Archaea

**DOI:** 10.1101/2021.12.15.472764

**Authors:** Jonathan Filée, Hubert F. Becker, Lucille Mellottee, Zhihui Li, Jean-Christophe Lambry, Ursula Liebl, Hannu Myllykallio

## Abstract

Little is known about the evolution and biosynthetic function of DNA precursor and the folate metabolism in the Asgard group of archaea. As Asgard occupy a key position in the archaeal and eukaryotic phylogenetic trees, we have exploited very recently emerged genome and metagenome sequence information to investigate these central metabolic pathways. Our genome-wide analyses revealed that the recently cultured Asgard archaeon *Candidatus* Prometheoarchaeum syntrophicum strain MK-D1 (*Psyn*) contains a complete folate-dependent network for the biosynthesis of DNA/RNA precursors, amino acids and syntrophic amino acid utilization. Altogether our experimental and computational data suggest that phylogenetic incongruences of functional folate-dependent enzymes from Asgard archaea reflect their persistent horizontal transmission from various bacterial groups, which has rewired the key metabolic reactions in an important and recently identified archaeal phylogenetic group. We also experimentally validated the functionality of the lateral gene transfer of *Psyn* thymidylate synthase ThyX. This enzyme uses bacterial-like folates efficiently and is inhibited by mycobacterial ThyX inhibitors. Our data raise the possibility that the thymidylate metabolism, required for *de novo* DNA synthesis, originated in bacteria and has been independently transferred to archaea and eukaryotes. In conclusion, our study has revealed that recent prevalent lateral gene transfer has markedly shaped the evolution of Asgard archaea by allowing them to adapt to specific ecological niches.

## Introduction

The canonical thymidylate synthase ThyA (EC 2.1.1.45) was once considered the only enzyme capable of catalyzing the *de novo* methylation of the essential DNA precursor dTMP (deoxythymidine 5’-monophosphate or thymidylate) from dUMP (deoxyuridine 5’-monophosphate). However, combined *in silico* and experimental approaches led to the discovery of a new metabolic pathway that operates in the methylation of DNA precursors in numerous microbial species ^1, 2, 3^, relying on a novel thymidylate synthase, ThyX (EC 2.1.1.148). No sequence or structural homology exists between ThyA (found in ≈ 65 % of microbial genomes) and ThyX flavoproteins (≈ 35 %)^4^. Differently from the homodimeric ThyA proteins, the active site of ThyX flavoenzymes, which accommodates dUMP, NADPH and the carbon donor methylene tetrahydrofolate (CH_2_H_4_folate, a vitamin B9 derivative), is located at the interface of three subunits of the homotetrameric protein complex^5^. Consequently, formation of the ThyX tetramer is necessary for catalytic activity.

In the unique reductive methylation reaction catalyzed by homodimeric ThyA, CH_2_H_4_folate functions both as a source of carbon (C_1_-carrier) and reducing equivalents^6^, thus leading to the formation of dihydrofolate (H_2_folate) as the oxidation product of tetrahydrofolate (H_4_folate) (Figure 1a, left panel). H_2_folate is subsequently reduced to tetrahydrofolate (H_4_folate) by dihydrofolate reductase (DHFR) FolA, as only reduced folate derivatives are functional in intermediary metabolism. Consequently, ThyA and FolA form a functionally coupled adaptive unit embedded within the bacterial folate metabolism network^7^. CH_2_H_4_folate is then reformed in a reversible reaction by serine hydroxymethyltransferase (SHMT) GlyA using H_4_folate and serine as substrates.

**Fig. 1.**
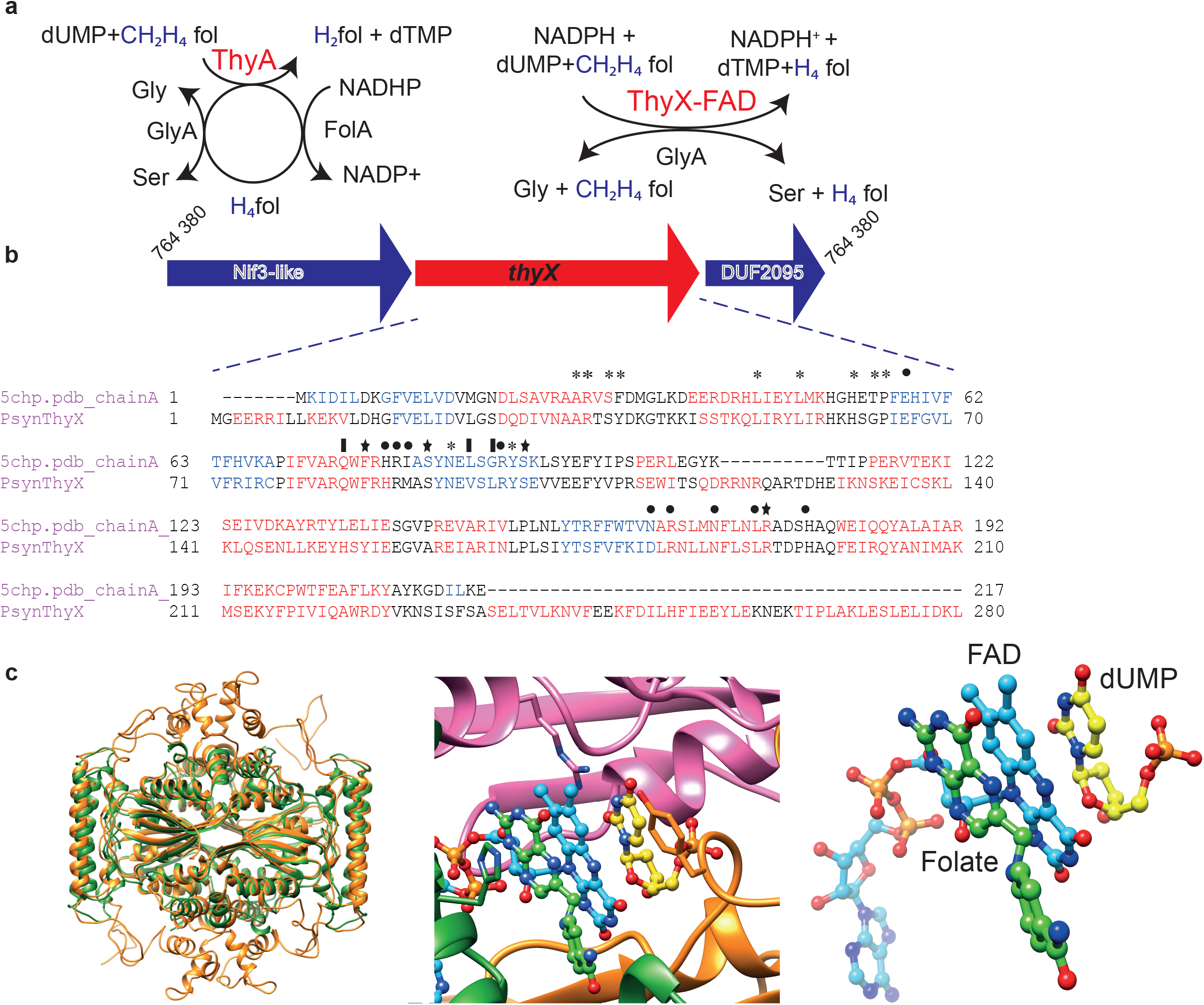
**a** ThyA- and ThyX-dependent folate cycles. ThyA catalyzes the methylation of dUMP to dTMP, leading to the formation of H2folate that is subsequently reduced by FolA (left panel). The flavoenzyme ThyX uses methylene from CH_2_H_4_folate and acquires the reducing hydride from NADPH. CH_2_H_4_folate functions only as a carbon source, resulting in H_4_folate at the end of the catalytic cycle (right panel). Serine hydroxymethyltransferase GlyA is universally present. **b** *Psyn thyX* is located between the nucleotides 765,745 and 764,847 on the genome of *Psyn* (NZ_CP042905, 4 427 796 bp). The *Psyn thyX* gene is surrounded by the *nif3* and DUF2095 genes. For details, see text. Structure-based sequence alignment of *Psyn* ThyX with *Thermotoga maritima* ThyX (corresponding to PDB structure 5CHP) is also shown. Functionally important residues are indicated above the alignment. Asterisks refer to folate binding, stars to nucleotide-binding and filled circles to flavin binding residues. **c** Structural model for the *Psyn* ThyX homotetramer (left panel) using PDB structures 1O26, 3N0B and 6J61 as templates. Superposition of the model with PDB structure 3GT9 allowed the addition of FAD (cyan), dUMP (yellow), and folate (green) molecules (middle panel) to the structural model. The substrate and co-factor configuration is highlighted in the right panel.

ThyX also uses CH_2_H_4_folate as a C_1_ carrier, but acquires the reducing hydride from NADPH and not from H_4_folate. ThyX catalyzes dTMP formation using flavin-mediated hydride transfer from pyridine nucleotides, which is required for reduction of the methylene group, thus directly resulting in the formation of H_4_folate (Fig. 1a, right panel). The ThyX reaction mechanism appears less catalytically efficient than that used by ThyA enzymes^8^, but maintains the folate in its reduced form (as H_4_folate) at the end of the catalytic cycle. Consequently, *thyX*-containing organisms do not have an absolute requirement for FolA in their thymidylate metabolism (Fig. 1a), explaining why the *folA* gene is frequently absent from *thyX*-containing organisms^9^. It remains unclear how two independent solutions for the synthesis of the central building block of DNA evolved. In addition, thymidylate synthase homologs are known to participate in biosynthesis of nucleoside antibiotics and modified nucleotides in bacteriophages^9^. Phylogenetic analyses have revealed the existence of frequent and multiple lateral gene transfers during the evolution of ThyA and ThyX, sometimes associated with non-homologous replacement^1, 10^. *ThyX* appears overrepresented in genome-reduced and/or slow-growing prokaryotes ^8, 11^, and changes in ThyX function of human microbiota associate with obesity-associated deficits in inhibitory control towards responses to stimuli (e.g. food)^12^.

Despite the essential role of thymidylate synthase for DNA synthesis, data and details on the experimental characterization of archaeal thymidylate synthases remain scarce. Consequently, the functionality of the archaeal thymidylate synthases has only been sporadically followed up to date. While the halophilic archaeon *Haloferax volcanii* contains the gene for canonical thymidylate synthase ThyA, in the closely related halophilic archaeon *Halobacterium salinarum* genetic evidence was provided for the presence of functional *thyX*^3^. Biochemically dTMP biosynthesis has been demonstrated in cell-free extracts of two different archaeal species^13, 14^: *Methanosarcina thermophila* and *Sulfolobus solfataricus* (now referred to as *Saccharolobus solfataricus*). Although the corresponding enzymes were not purified or identified in this study, dTMP formation was detected from dUMP using externally added isotope-labeled formaldehyde and either chemically modified folates present in the cell extract (tetrahydrosarcinapterin, *M. thermophila*) or added synthetic fragments of sulfopterin (*S. solfataricus*). Indeed, biochemical studies have indicated the presence of at least six distinct chemically modified, but thermodynamically similar, C_1_ carriers that function in archaeal central biosynthetic networks and/or energy-yielding reactions^15^. These chemical modifications differ among various archaeal species and likely replace chemically distinct tetrahydrofolate derivatives in many archaea. Recent phylogenetic studies have suggested an archaeal origin for the energy-producing Wood-Ljungdahl pathway that is dependent on tetrahydromethanopterin (H_4_MPT), frequently found in methanogenic archaea^16^. This archaeal pathway may have contributed to the early origin of methanogenesis and the emergence of the use of *e*.*g*. reduced one-carbon compounds as carbon source (methylotrophy) in bacteria.

An archaeal superphylum called Asgard was recently discovered that includes the closest known archaeal relatives of eukaryotes^17, 18, 19^. This striking discovery has solidified the two-domains tree of life hypothesis in which Eukaryotes have emerged from within the archaeal tree^20^. Even if the true identity of the archaeal ancestors of Eukaryotes is still being debated, Asgard archaea occupy a pivotal position in the archaeal/eukaryotic phylogenetic trees^17, 19^. Thus, studying the thymidylate and folate metabolism in the Asgard group is of particular interest to better understand the origins and the evolution of these essential pathways participating in DNA, RNA, and protein synthesis, as well as in catabolic reactions. This is also underlined by the fact that pioneering genomic studies on Lokiarchaeota suggested the absence of biosynthetic capacity for tetrahydromethanopterin and tetrahydrofolate co-factors, but the presence of some folate-dependent enzymes implicated in amino acid utilization^18^.

In this study, we have exploited the recent increase in Asgard metagenome and genome sequence information to investigate thymidylate synthases and folate-dependent metabolic networks in more than 140 complete or nearly complete Asgard genomes. Our sequence similarity searches revealed a high level of similarity between the protein sequences of bacterial and Asgard folate-dependent enzymes, including thymidylate synthases. Our genomic and phylogenetic analyses indicated that Asgard archaea have ‘hijacked’ at several independent occasions bacterial folate-dependent enzymes and pathways to support their central metabolism. Detailed experimental analyses revealed that the protein encoded by *thyX* from the recently cultured Asgard archaeon *Candidatus* Prometheoarchaeum syntrophicum strain MK-D1^18^ (further referred to as *Psyn*) is fully functional as thymidylate synthase in bacterial cells and efficiently interacts with bacterial folates. In conclusion, our combined experimental and computational data suggest that the patchy phylogenetic distribution and phylogenetic incongruences of functional folate-dependent enzymes from Asgard archaea reflect their independent horizontal transmission from various bacterial groups. These lateral (horizontal) gene transfer (LGT) events have potentially rewired the key metabolic reactions in an important archaeal phylogenetic group.

## Results

### Identification of a ThyX orthologue in *Candidatus Prometheoarchaeum syntrophicum* strain MK-D1 (*Psyn*)

As an initial approach to investigate the Asgard archaeal thymidylate and C_1_ metabolism, we searched for *thyX* and *thyA* homologs in sequence databases for archaeal genomes or metagenome-assembled genomes (MAGs). These studies led to the identification of both *thyX* and *thyA* sequences in the genomes of the understudied and diverse group of archaea. Interestingly, our similarity searches identified a ThyX orthologue in *Psyn*, which is up to 51.44% identical to bacterial ThyX sequences at the protein sequence level (e.g. *Zixibacteria*, e-value ≈ 10^−78^ or *Calditrichaeota* bacterium, e-value ≈ 10^−76^). The observed e-values are very low and are adjusted to the large sequence database size. Therefore, the quality of the observed hits is very high. As *Psyn* is the only known example of *Asgard* archaea that can be cultivated and its genome, lacking *thyA*, is completely sequenced, we concentrated our efforts on this archaeal ThyX.

Transcriptome analyses of RNA extracted from enriched cultures of *Psyn* indicated that *thyX* from this strain is expressed [Reads Per Kilobase of transcript, per Million mapped reads (RPKM) value of 225.37 using the data set from the sequence read archive (SRA) DRR199588]. This gene encodes a protein with a predicted molecular mass of 35, 294 Da. Its genomic environment (Fig. 1b and Supplementary Figure 1) comprises an upstream gene coding for a Nif3-like protein with a length of 259 residues (29,026 Da) and a downstream gene encoding a domain of unknown function DUF2095. Physical association of *Psyn thyX* with the Nif3 family encoding gene is of interest, as this family of proteins may correspond to GTP cyclohydrolase 1 type 2, which converts GTP to dihydroneopterin triphosphate and may function in folic acid synthesis. The structure-based sequence alignment of *Psyn* ThyX with *Thermotoga maritima* ThyX (PDB structure 5CHP) predicts the conservation of functionally important residues involved in folate-, nucleotide- and flavin-binding (Fig. 1b). The marked degree of sequence similarity allowed the construction of a high-quality structural model for *Psyn* ThyX indicating its functional significance as thymidylate synthase and nucleotide and folate binding protein. More specifically, we constructed a model of the ThyX homotetramer based upon PDB structures 1O26, 3N0B and 6J61 as templates and using the protein structure modeling program Modeller^21^ (Fig. 1c, left panel). Structural superposition of the model with PDB structure 3GT9 using the Chimera software^22^ allowed the addition of FAD, dUMP and folate molecules into the active site of the model (Fig. 1c, middle panel). Importantly, the model suggests the highly plausible transfer of 1C units via the N5 atom of the FAD cofactor to the accepting carbon of dUMP (Fig. 1c, right panel).

### thyX and thyA are present in Asgard archaea

As our sequence similarity searches suggested a close link between *Psyn* and bacterial ThyX sequences, we obtained more detailed insight into the thymidylate synthase gene distribution and phylogeny in Asgard archaea. In particular, in addition to the complete genome of cultivated *Psyn*, we analyzed more than 140 MAGs of uncultivated Asgard archaea (Fig. 2). Our analyses revealed that *thyX* is present in only a small subset of lineages with a patchy gene distribution. We found *thyX* genes to be present in 33 out of a total of 141 Asgard (meta)genomes: 6/52 in the Loki lineage, 6/16 in Heimdall, 0/5 in Hel, 1/1 in Odin, 6/38 in Thor, 1/1 in Wukong, 1/2 in Borr, 1/12 in Hod, 3/3 in Gerd, 7/10 in Hermod, 0/1 in Kari and 2/2 in Baldr. In contrast, *thyA* genes are widely distributed among Asgard archaea: 93 out of a total of 141 (36/52 for Loki, 13/16 for Heimdall, 5/5 for Hel, 1/1 for Odin, 31/38 for Thor, 0/1 for Kari, 1/2 for Wukong, 1/2 for Borr, 3/12 for Hod, 0/3 for Gerd, 1/10 for Hermod, 1/2 for Baldr). Approximately twenty Asgard MAGs encode for both *thyX* and *thyA* genes, although *thyX* genes appear often partial or fragmented in this latter case. In a few cases, both *thyX* and *thyA* appear to be absent, likely reflecting the fact that some of the analysed MAGs correspond to partially assembled genomes.

**Fig. 2.**
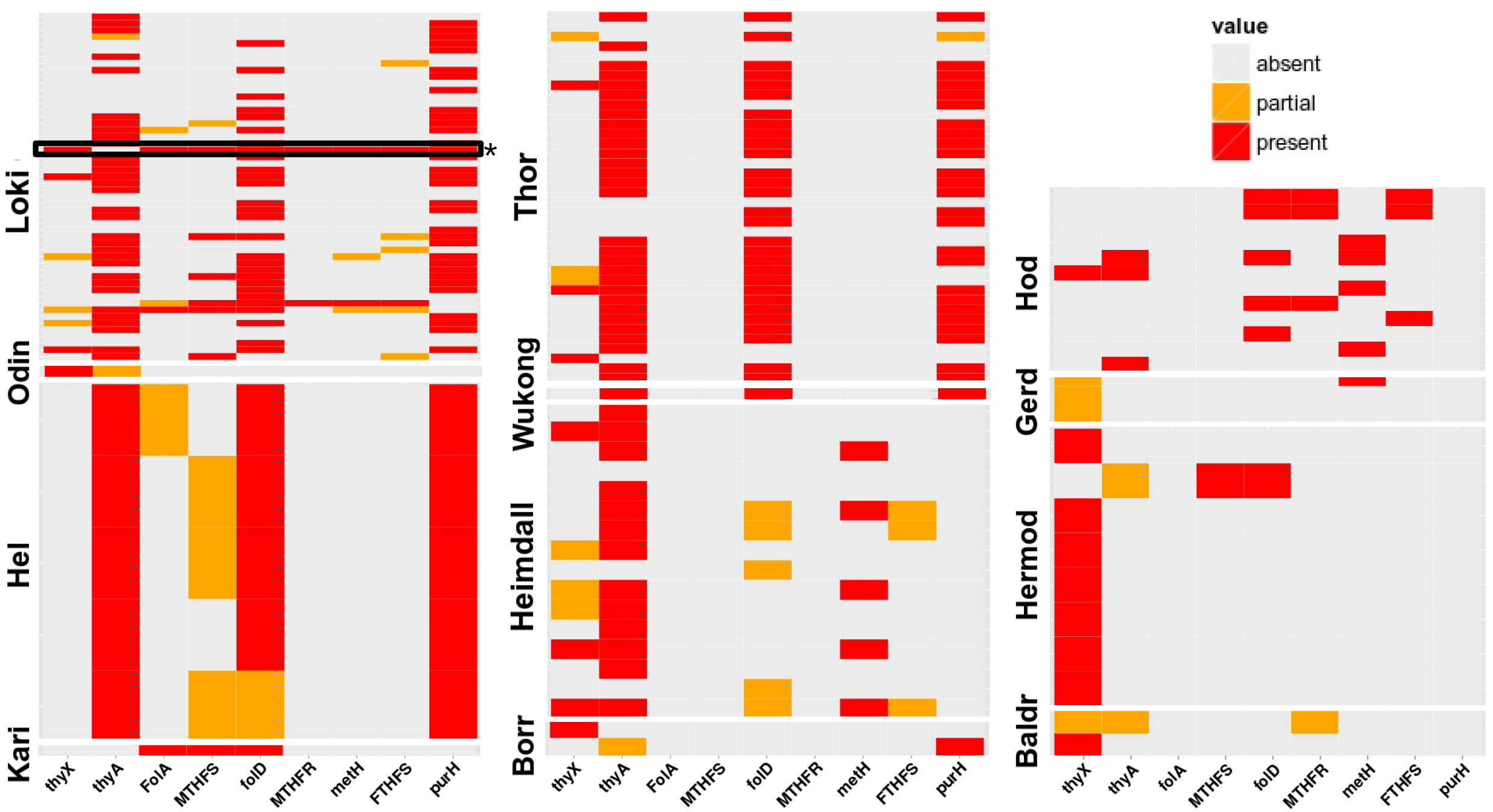
Heatmap for phyletic distribution of different folate-dependent enzymes in different phyla of Asgard archaea. The red color indicates the presence of the entire gene, orange designates a partial copy. *Psyn* is indicated with an asterisk and its gene distribution is framed in black. The (meta)genomic assembly and gene names are in supplementary table 1 (sheet “distribution of folate genes”).

### Automated prediction of gene transfers between *Psyn* and bacteria

Considering the high level of sequence similarity of *Psyn* ThyX and bacterial orthologs, we estimated the possible extent of gene transfer between bacteria and *Psyn* using HGTector. This automated method is well suited for discovering potential recent gene transfers^23^ by analyzing the hit distribution patterns from the similarity searches. The method performs a systematic analysis to detect genes that do not appear to have a typical vertical history of the analysed organisms but instead have a putative horizontal origin. This approach efficiently detects atypical genes that are highly likely to be horizontal gene transfers (HGT). As this method uses all sequence data available in the GenBank, it allows exhaustive detection of potential recent gene transfers.

Using HGTector, we predicted 149 HGT events [supplementary table 1 (sheet “potential gene transfers”)], including *Psyn thyX*, when all the ≈ 4000 *Psyn* predicted protein-coding genes were analyzed (Fig. 3). In this plot, each dot represents one gene, and likely horizontally transferred genes are indicated in yellow. As depicted in Fig. 3, many *Psyn* folate-dependent genes (purple legends in Fig. 3a, more detailed figure with annotations is available as supplementary Fig. 3) were also likely transferred from bacteria. The BlastKOALA annotation^24^ of the potentially transferred genes indicates their functional distribution in amino acid, nucleotide, and carbohydrate metabolism (Fig. 3b and supplementary table 1). This is of interest considering syntrophic amino acid utilization of *Psyn*^18^.

**Fig. 3.**
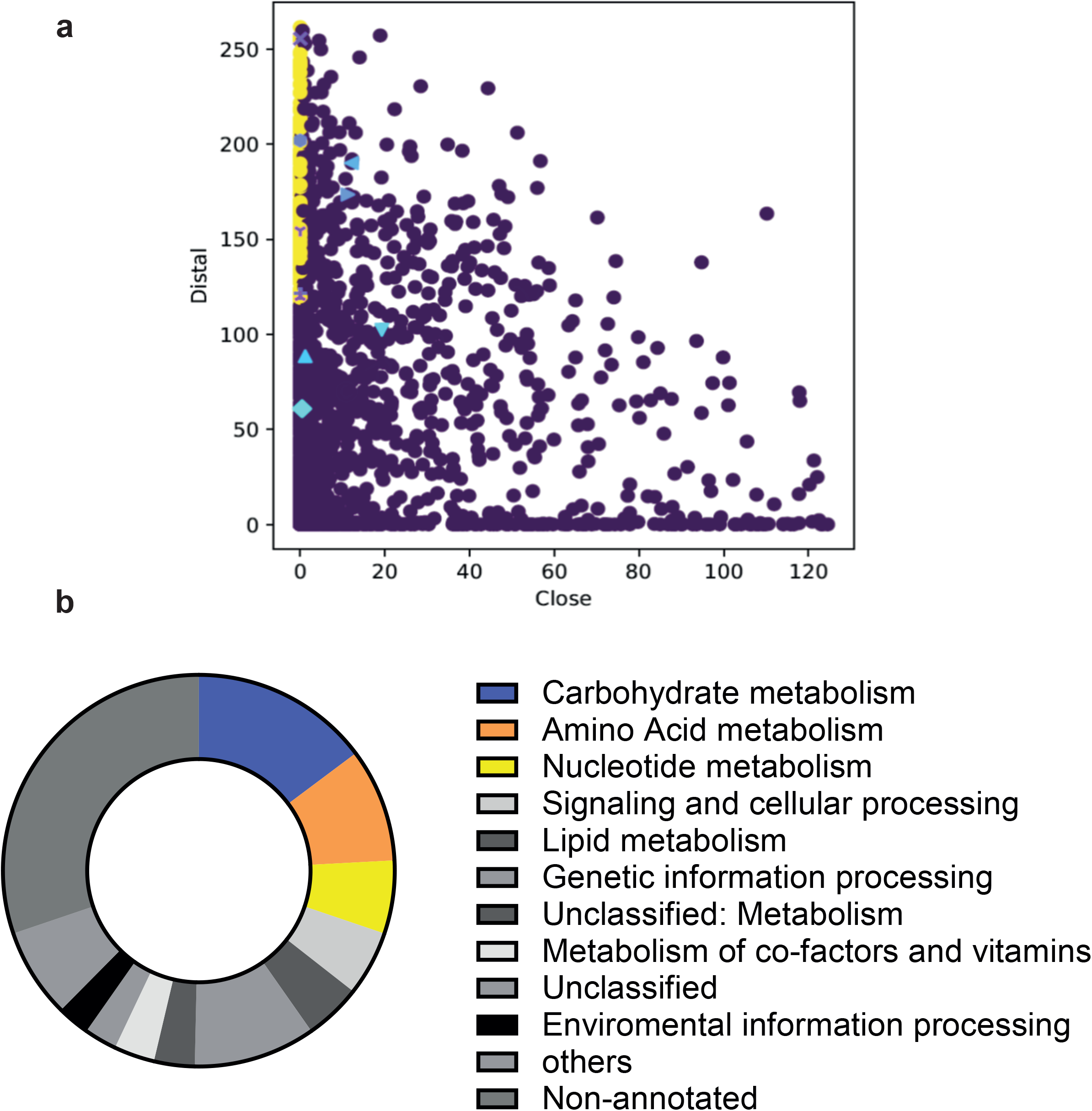
Automated prediction of gene transfers between *Psyn* and bacteria. **a** Plot indicating the distributions of “close” and “distal” scores revealed by similarity searches. The scatter plot was created by the HGTector. Potentially transferred genes (n=149) are indicated in yellow, whereas the other coloured labels refer to *Psyn* folate-dependent enzymes (for annotation, see supplementary Figs. 2 and 3). **b** Distribution of the functional groups for potentially transferred *Psyn* genes obtained using the BlastKOALA annotation^24^.

### *thyX* was transferred from bacteria to diverse Asgard archaea

To better understand the origin and evolution of the thymidylate synthase genes in Asgard archaea, we have reconstructed their phylogenies. Note that Asgard sequences are indicated in red, other archaea in blue, bacteria in black and Eukarya in green. The ThyX phylogeny (Fig. 4 and supplementary Fig. 4 for a detailed version of the tree with the taxon names) shows that Asgard sequences are deeply polyphyletic; they are scattered into several monophyletic groups. Four of them are related to different bacterial groups and appear distantly related to other archaeal sequences. The *Psyn* ThyX clusters along with some, but not all, Loki and Heimdall sequences, together with bacteria belonging to the *Deinococcus*/*Thermus* group, as well as with several alpha-proteobacteria. Additional *Heimdall* and *Borrarchaeota* sequences appear scattered in this tree with varying affinities for diverse bacterial lineages. The three remaining Asgard groups, including all of the thorarchaeal sequences, are related to other archaeal sequences, but do not form a monophyletic cluster, indicating possible lateral gene transfers among archaeal species. The only eukaryotic ThyX sequence from *Dictyostelium* branches with alpha-proteobacteria, suggesting a transfer from bacteria to eukaryotes. This phylogenetic analysis supports the existence of multiple and independent lateral (horizontal) gene transfers of the *thyX* genes from highly diverse bacteria into Asgard genomes.

**Fig. 4.**
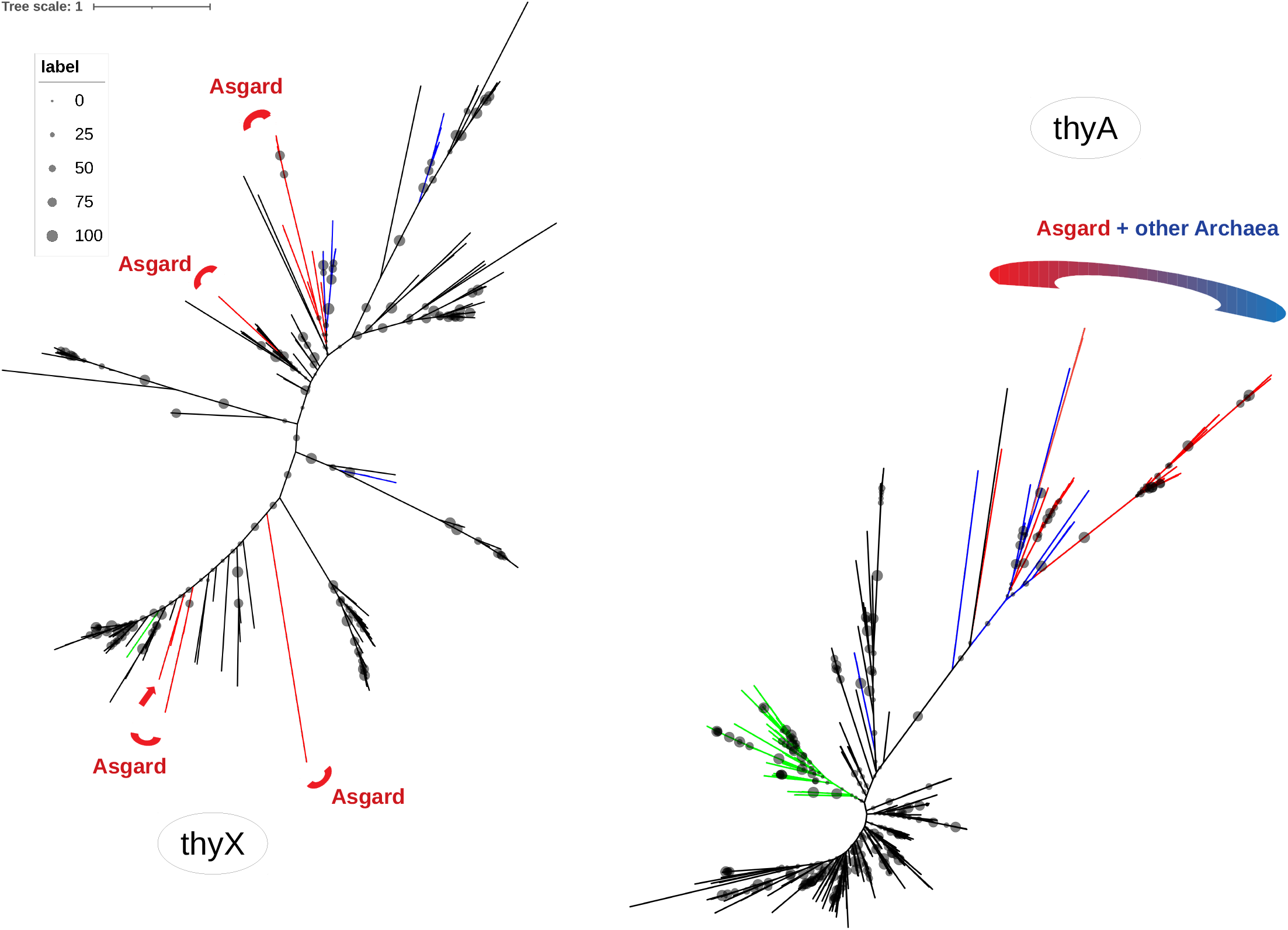
Unrooted Maximum likelihood phylogenetic tree of ThyX (198 sequences, alignment of 157 amino acids with the LG+G+I phylogenetic model) and ThyA (390 sequences, alignment of 121 amino acids with the LG+G+I model) with a focus on Asgard proteins. The obtained ThyX tree indicates multiple lateral gene transfers from bacteria to *Psyn* and other Asgard genomes. By contrast, the ThyA tree refutes the occurrence of recent and multiple lateral gene transfers from bacteria to Asgard genomes. Asgard sequences are indicated in red, other archaea in blue, bacteria in black, and eukarya in green. The scale bar indicates the average number of substitutions per site. Bootstrap values are proportional to the size of the circles on the branches.. The complete trees are available in the supplementary data.

### ThyA corresponds to the ancestral thymidylate synthase in *Asgard* archaea

In contrast to ThyX, the ThyA phylogeny (Fig. 4 and supplementary Fig. 5 for a detailed version of the tree with the taxon names) indicates that Asgard sequences form a monophyletic group within various archaea. The ThyA sequences of the Asgard archaea are distributed into four clusters that appear related to diverse methanogenic archaea. This pattern is incompatible with the occurrence of multiple, recent, and independent lateral gene transfers from bacteria as observed above with ThyX (Fig. 4). Finally, the Asgard ThyA sequences appear distant from bacterial and eukaryotic sequences, suggesting that the ancestors of the Asgard archaea contained ThyA. However, this tree topology is also compatible with the more complex scenarios of *i)* the Asgard ancestor inheriting its *thyA* gene from a bacterium, followed by subsequent gene transfers to diverse methanogens, or *ii)* initial *thyA* transfers to methanogenic archaea and subsequent ones to the Asgard ancestor.

### *Psyn thyX* is functional in *Escherichia coli*

Next, we investigated whether *Psyn thyX* functionally complements growth defects of an *E. coli* strain specifically impaired in thymidylate synthase activity. Towards this goal we designed and constructed a synthetic plasmid, pTwist-Psyn-ThyX, where the transcription of *Psyn thyX* is under control of a synthetic T5 promoter, carrying the *lac* operator (*lacO*) sequences. The plasmid also contains an ampicillin resistance marker and a p15A replication origin, as well as a λ t0 terminator, located after the *Psyn thyX* gene (Fig. 5a). As the T5 promoter is recognized by native *E. coli* RNA polymerase, this plasmid expresses an N-terminal His-tagged *Psyn* ThyX protein in any *E. coli* strain.

**Fig. 5.**
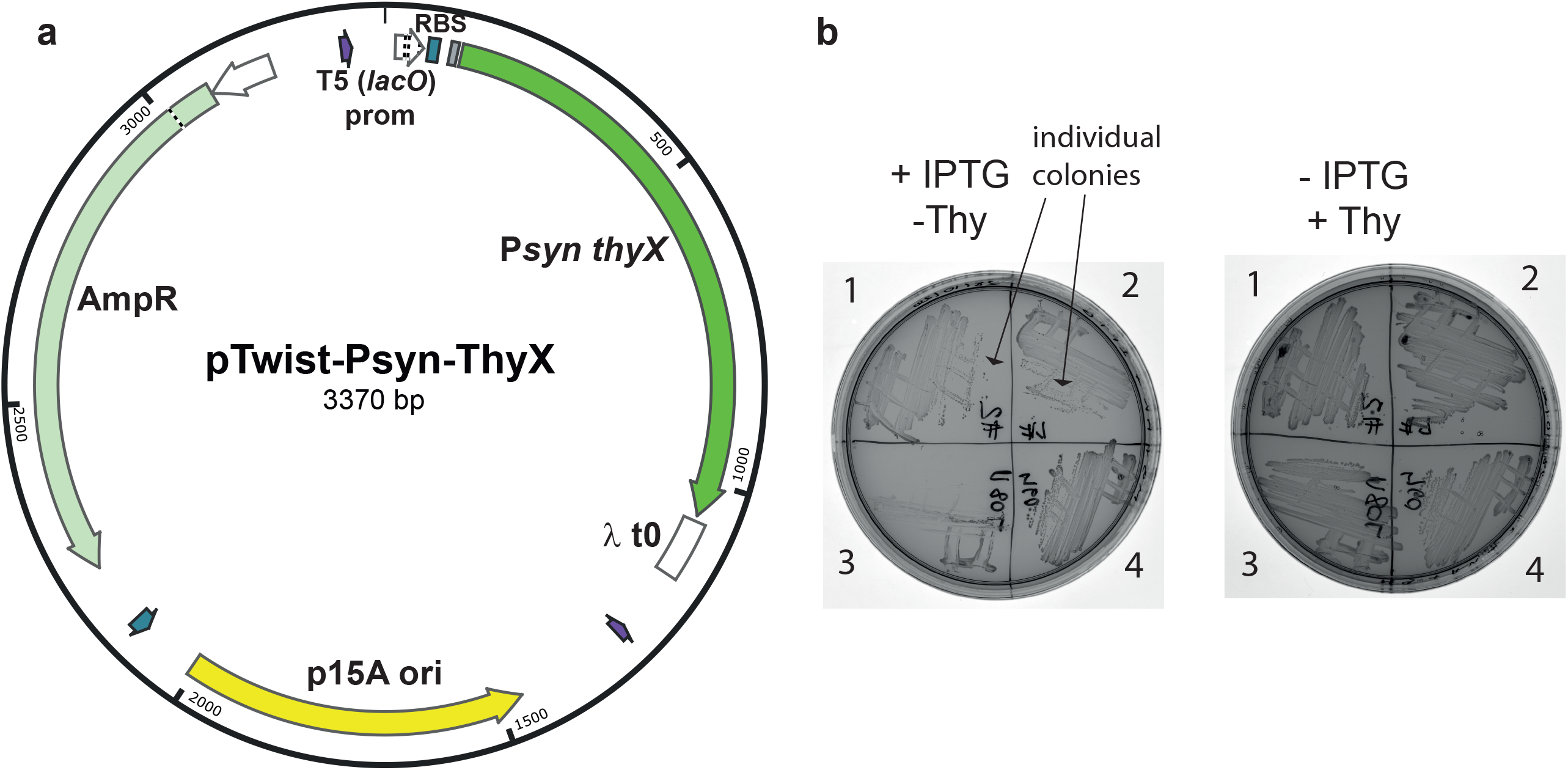
**a** Map for the synthetic plasmid pTwist-Psyn-ThyX. Transcription of the *Psyn thyX* gene is under the control of a synthetic T5 promoter carrying *lacO*. An ampicillin resistance marker and a p15A replication origin are integrated, as well as a lambda *t0* terminator. **b** Interkingdom complementation assay using *Psyn thyx*. The ability of *Psyn thyX* to permit thymidine-independent growth of *E. coli* thymidine-auxotroph FE013^8^ (Δ*thyA, lacI*) was scored after three days at 37°C in the presence of 1mM IPTG using minimal M9 medium. 1 and 2 correspond to two independent clones of *Psyn thyX*/FE013 whereas 3 and 4 are negative (no insert) and positive (*C. trachomatics thyX*) controls, respectively. The plate on the left contains IPTG, but no thymidine, whereas the plate on the right contains thymidine, but no IPTG.

The ability of *Psyn thyX* to permit thymidine-independent growth of the *E. coli* thymidine-auxotroph strain FE013^8^ (Δ*thyA::aphA3*, derived from wild type MG1655) was scored after three days at room temperature or 37°C in the presence of 1mM IPTG using either minimal M9 or thymidine-deprived rich medium L^+^ (see the methods section). Fig. 5b shows the formation of individual colonies of *Psyn thyX*/FE013 in the absence of thymidine in the presence of IPTG and appropriate antibiotics (plate on left, streaks 1 and 2). Under these thymidine limiting conditions, the strain carrying the control plasmid lacking the insert did not form individual colonies (streak 3, Fig. 5b). Altogether the results of this inter-kingdom complementation experiment indicate that this archaeal ThyX protein is fully functional in a bacterial strain and must use *E. coli* folate derivatives for *de novo* thymidylate synthesis.

### Enzymatic activity of *Psyn*ThyX

The recombinant histidine-tagged *Psyn*ThyX was produced in soluble form and efficiently purified by one-step affinity chromatography (Figure 6a). Western blot analysis using an anti-Histidine-Tag monoclonal antibody identified that revealed a single band with an apparent molecular mass of ∼37 kDa (Figure 6b). Thymidylate synthases ThyX are dUMP-dependent NADPH oxidases^25, 26^, differently from thymidylate synthase ThyA. The formation of dTMP catalyzed by ThyX enzymes involves two half-reactions, oxidation and transfer of the methylene group from CH_2_H_4_Folate to dUMP, to form the final product dTMP. Using a spectrophotometric biochemical assay, we found that *Psyn* ThyX catalyzes the NADPH oxidation to NADP^+^ only in the presence of dUMP (Figure 6c). The oxidation of NADPH was assayed with 0.4μM *Psyn*ThyX using saturating amounts of FAD co-factor (50 μM) and the substrates dUMP (20 μM) and NADPH (750 μM). The specific activity of *Psyn*ThyX (0.030 UI.mg-1) is somewhat lower than that of *Mycobacterium tuberculosis* ThyX (0.044 UI.mg-1) or *Paramecium bursaria chlorella virus-1* (PBCV-1) ThyX (0.043 UI.mg-1). However, this level of activity is sufficient for genetic complementation (Fig. 5b). Under these experimental conditions, the hyperbolic saturation curves show a nanomolar affinity of the nucleotide substrate for the enzyme [K_m, dUMP_ = 235 ± 35 nM (Figure 6d)].

**Fig. 6.**
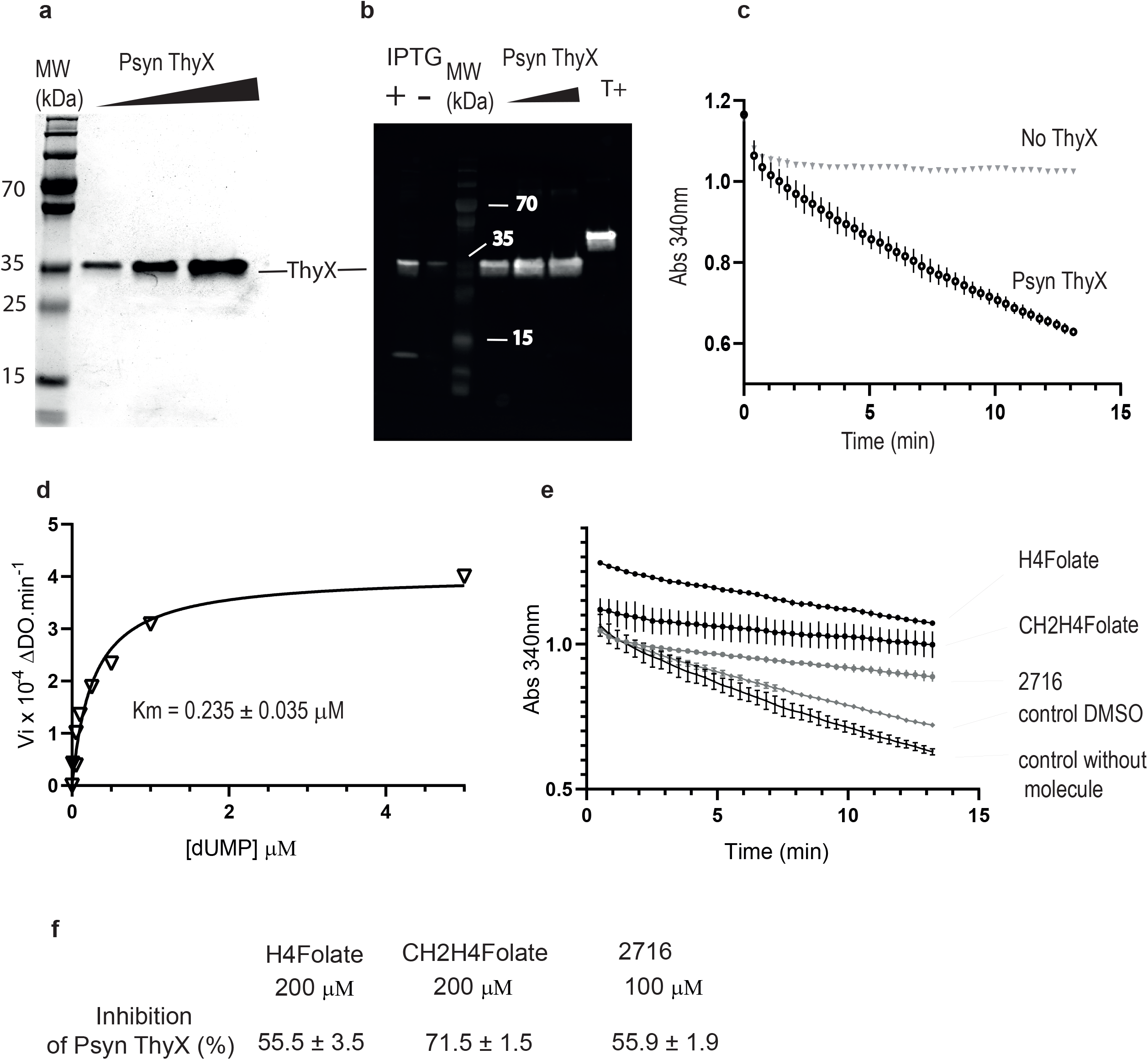
Biochemical characterization and inhibition analyses of *Psyn* ThyX. **a** 12% TGX™ PAGE of *Psyn* ThyX purified by one-step affinity chromatography. 0.5, 1, and 2 micrograms of purified protein were loaded from the left to the right. MW: molecular marker. **b** Western immunoblot of IPTG-induced and non-induced whole cell lysates of *E. coli* FE013 using anti-His monoclonal antibodies. A single band with an apparent molecular mass of ∼37 kDa is revealed. T^+^: control lane (*Pyrococcus abyssi* RNA ligase). **c** Representative NADPH oxidation activity curves of *Psyn* ThyX as measured following the absorption change at 340 nm. The control without enzyme is also shown. Assay mixtures contained 0.4 μM enzyme, 50 μM FAD, 20 μM dUMP, and 750 μM NADPH. **d** Oxidation rate of NADPH under saturating dUMP concentrations. The affinity of dUMP for *Psyn* ThyX was determined from the hyperbolic saturation curve. **e** Evaluation of inhibiting properties of H_4_folate (200 μM), CH_2_H_4_folate (200 μM), and the folate analog ‘2716’ (100 μM) ^23, 24^. Molecule ‘2716’ is solubilized in DMSO. Controls: without molecule and in presence of 1% DMSO. **f** Percentage of inhibition calculated for the three folate derivatives or inhibitors. Standard deviations (S.D) of three independent measurements are also shown.

### Inhibition of *Psyn*ThyX by folate analogs

Our genetic complementation tests indicate that *Psyn* ThyX must use bacterial folates for thymidylate synthesis. To provide additional support for this notion, we further characterized the *Psyn*ThyX enzymatic activity by performing kinetics measurements in the presence of molecules that bind to the folate binding pocket of bacterial ThyX proteins. These inhibitory studies of *Psyn* ThyX were performed using H_4_folate (reaction product), CH_2_H_4_folate (substrate), and the tight-binding *Mycobacterium tuberculosis* ThyX inhibitor 2716^27, 28^, which all inhibit the NADPH oxidase activity of bacterial ThyX. This analysis revealed that the three bacterial folate analogs tested substantially inhibit *Psyn* ThyX activity compared to an assay without the addition of any molecule (Fig. 6e). Note that the molecule 2716, a potent inhibitor of *Mycobacterium tuberculosis* ThyX, was solubilized in DMSO and results need to be compared to the control condition in presence of 1% DMSO. The three folate analogs presented a percentage of inhibition of *Psyn*ThyX over 50% with high reproducibility (Figure 6f). Altogether our genetic (Fig. 5) and biochemical (Fig. 6) data indicate that bacterial folate-like molecules are efficiently utilized by archaeal *Psyn* ThyX for *de novo* synthesis.

### Distribution of the folate-mediated one-carbon metabolism enzymes in *Asgard* archaea

As THF derivatives are important biological cofactors that play a wide role in the biosynthesis of DNA, RNA, and proteins, our observations obtained using *PsynThyX* prompted us to investigate in more detail *Psyn* folate-dependent biosynthetic networks in genomes and metagenomes of Asgard archaea.

Our analyses indicated the sporadic presence of many C_1_ generating/transferring and folate-interconverting enzymes in Asgard archaea (Fig. 2). In addition to *thyX* and *thyA* genes, we have analyzed the distribution of *folA*, 5-formyltetrahydrofolate cyclo-ligase (MTHFS), *folD*, methylenetetrahydrofolate reductase (MTHFR), *metH*, formate-tetrahydrofolate ligase (FTHFS), and *purH* genes in the *Psyn* genome and other Asgard metagenomic assemblies (Fig. 2 and supplementary table 1). Many of these were predicted as likely transfer events by the HGTector (Fig. 3 and supplementary Fig. 3). ThyA appears the most universal folate-dependent *Asgard* enzyme, whereas the majority of the other folate-related genes have a very sporadic distribution. Nevertheless, the *folD* and *purH* genes are frequently found in *Asgard* archaea except for the *Heimdall* and *Odinarchaeota*, where they are absent or partially deleted.

The phylogenies of FTHFS and PurH show that a substantial fraction of the Asgard sequences are grouped with other Archaea (Fig. 7 and supplementary Figs 6-11 for a detailed version of the trees with the taxon names). However, several Asgard sequences are only distantly related to other archaeal sequences and located close to bacterial ones. The Asgard FolA, MTHFS, MTHFR, and MetH sequences appear all related to bacterial sequences, indicating additional multiple and independent events of lateral interkingdom acquisition. By contrast, the FolD sequences (supplementary figure 11), which are widely distributed in Asgard genomes, form a monophyletic group with few bacterial sequences. This supports the idea that FolD may be the only ancestral folate-dependent enzyme in *Asgard* archaea, without obvious evidence of LGT. Therefore, with the possible exception of FolD, the phylogenetic trees of the folate-related enzymes analyzed above strongly support multiple LGT events from Bacteria (Fig. 7), as Asgard sequences appear as polyphyletic groups that are scattered into the bacterial subtrees in most of the obtained phylogenies.

**Fig. 7.**
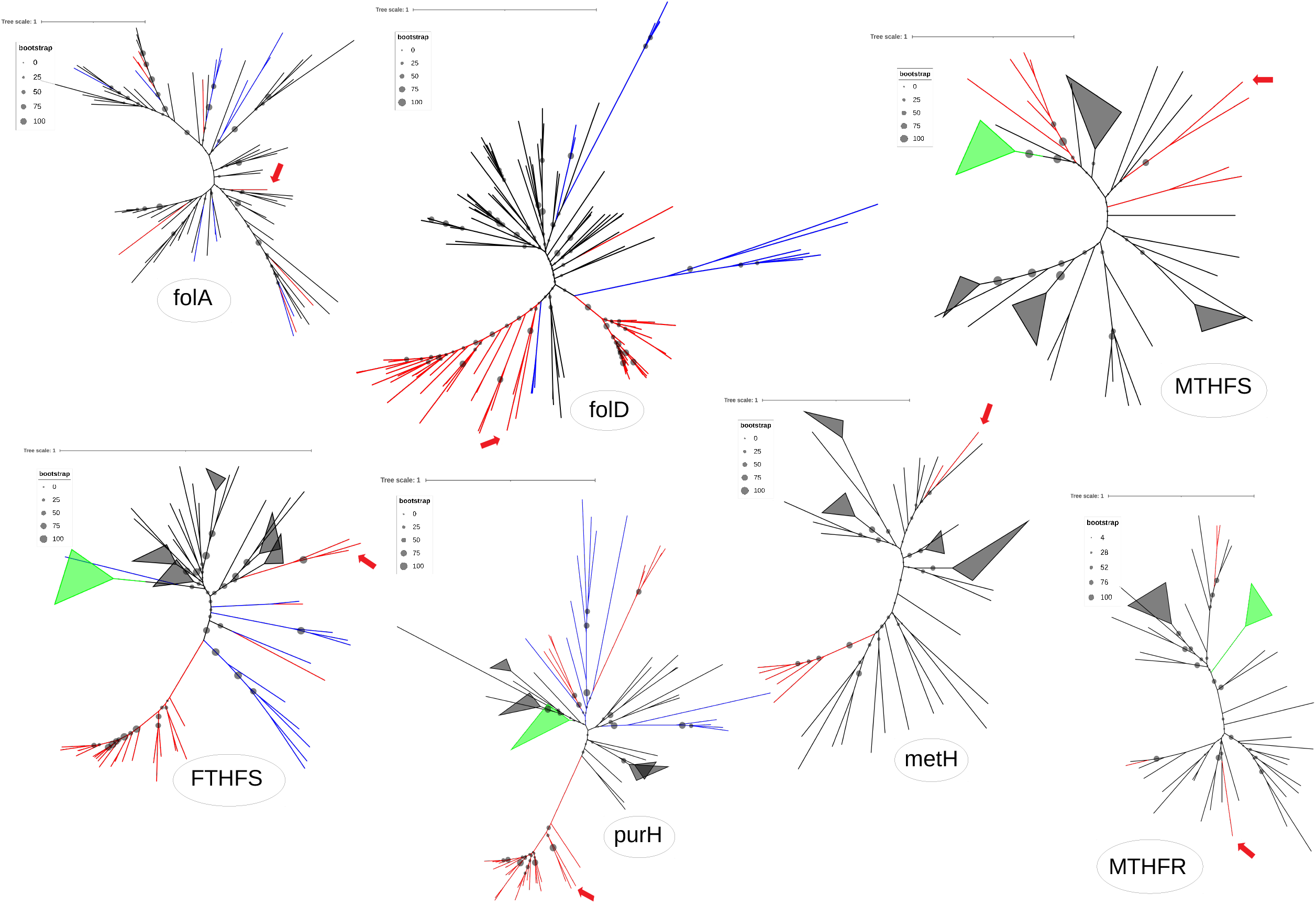
Unrooted Maximum likelihood phylogenetic tree of the folate-related proteins : FolA (83 sequences, alignment of 153 amino acids with the CpREV+G phylogenetic model), FolD (130 sequences, alignment of 244 amino acids with the BLOSUM62+G model), MTHFS (56 sequences, alignment of 149 amino acids with the LG+G model), FTHFS (79 sequences, alignment of 509 amino acids with the BLOSUM62+G model), PurH (94 sequences, alignment of 437 amino acids with the BLOSUM62+G model), MetH (47 sequences, alignment of 131 amino acids with the BLOSUM62+G model) and MTHFR (53 sequences, alignment of 280 amino acids with the BLOSUM62+G model) with a focus on Asgard proteins. The obtained trees indicate multiple lateral gene transfers from bacteria to *Psyn* and other Asgard genomes for most folate genes. Asgard FolD sequences form a monophyletic group with the other Archaeal sequences refuting gene transfer occurrences. Asgard sequences are indicated in red and *Psyn* with red arrows, other archaea are in blue, bacteria in black, and Eukarya in green. The scale bar indicates the average number of substitutions per site. Bootstrap values are proportional to the size of the circles on the branches. The complete trees are available in the supplementary data.

### Reconstruction of the complete folate-mediated one-carbon metabolism network in *Psyn*

*Psyn* is unique among *Asgard* archaea, as this cultivated species contains, based upon the MAGs analyzed, a remarkable number of predicted folate-dependent enzymes (Fig. 2 and supplementary table 1). This prompted us to perform a detailed reconstruction of enzymatic reactions participating in an interdependent reaction network required for chemically activating and transferring one-carbon (C_1_) units in *Psyn*. Our automated genome-wide analyses, together with highly sensitive manual similarity searches, revealed a high metabolic potential of *Psyn* to use folate derivates for de *novo* synthesis of not only thymidylate but also of inosine-5’-monophosphate (IMP), and remethylation of homocysteine to methionine (Fig. 8a). Moreover, we determined that both carbon dioxide (at the level of 10-formyl THF) and serine (at the level of CH_2_H_4_ THF) could provide a feasible source for one-carbon units entering the folate-mediated metabolic network in this species. The complete glycine cleavage system and its folate-binding component GcvT also provide an alternative source of CH_2_H_4_folate from glycine. According to our metabolic reconstruction studies, the folate network moreover participates in histidine catabolism by converting histidine to glutamate and 5-formyl THF. Many of these predictions are well supported by the strong association of genes encoding folate-dependent enzymes with the physically and functionally linked genes of, among others, glycine cleavage or histidine catabolism (Fig. 8b).

**Fig. 8.**
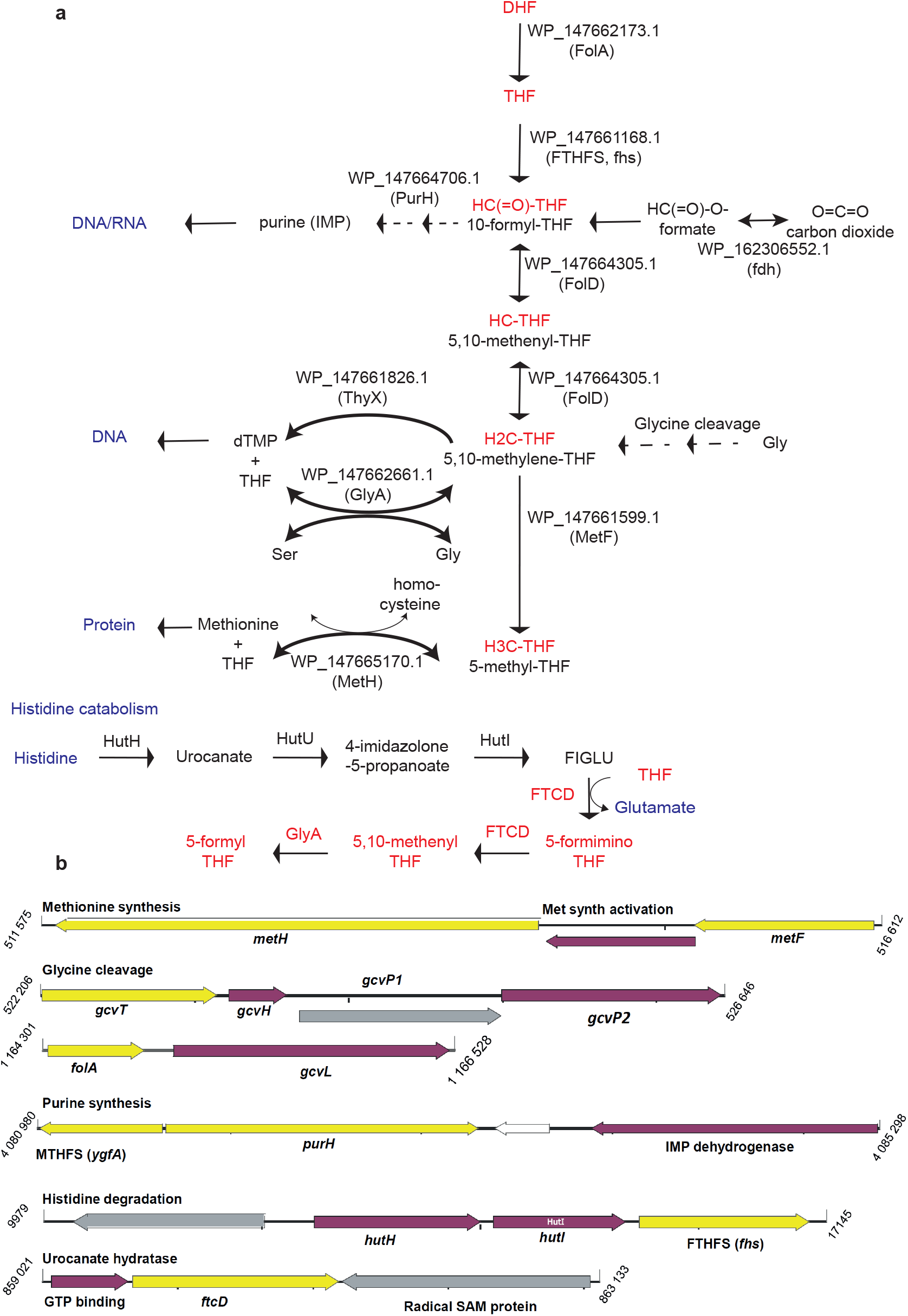
**a** Automated and manual procedures were used for metabolic reconstruction of the complete *Psyn* folate-dependent metabolism. Abbreviations not explained in the figure are DHF, dihydrofolate; THF, tetrahydrofolate; IMP, inosine monophosphate. **b** Functional connections between the folate-dependent *Psyn* enzymes (indicated in yellow) as suggested by the genomic context analyses. Purple and gray refer to genes functionally linked with the folate-dependent enzymes. For additional discussion, see text.

## Discussion

We initiated this study to obtain experimental and evolutionary information on thymidylate synthases in Asgard archaea. This led to the first experimental characterization of the archaeal flavin-dependent thymidylate synthase ThyX^2^ from *Psyn*. Our genetic and biochemical experiments reveal that this enzyme functions as a robust thymidylate synthase in bacterial cells (Fig. 5). Our biochemical results further indicate that *Psyn* ThyX catalyzes dUMP dependent NADPH oxidation, which is significantly inhibited by bacterial folate analogues (Fig. 6), as has been previously shown for bacterial or viral ThyX proteins (for a recent review, see^9^). Considering that archaea are widely thought to use C_1_ carriers chemically distinct from their bacterial counterparts, our biochemical observations are unexpected.

Our phylogenetic studies indicated for the first time that Asgard archaeal *Psyn* ThyX originated in Bacteria (Fig. 4). An alternative explanation could be the possible contamination of the *Psyn* genome assembly with foreign DNA. To exclude this possibility, we analysed the genomic environment of the *Psyn thyX* gene (Fig. 1 and supplementary Figure 1). Genes surrounding *Psyn thyX* are unambiguously of archaeal origin, as almost all of the 18 genes upstream or downstream have their first BLAST hit matching with an archaeon. Moreover, 14 of these genes have more than 75% of their top100 BLAST hits belonging to archaea. This analysis indicates that the *Psyn thyX* gene is inserted into an archaeal-like genomic context and is not the result of contamination or an assembly artifact. Thus, this gene represents a *bona fide* thymidylate synthase gene acquired laterally from bacteria. Taken together, our phyletic and phylogenetic analyses of thymidylate synthase genes in Asgard archaea reveal a complex evolutionary history involving many lateral gene transfers from bacteria and subsequent homologous and non-homologous replacements. The large distribution of *thyA* in Asgard metagenomes and the phylogenetic clustering of thymidylate synthase genes (Figs 4 and 5) along with the other archaeal sequences suggest that *thyA* is the ancestral thymidylate synthase, likely in Asgard and possibly in other archaea. This indicates that in *Psyn* the *thyX* gene likely replaced the ancestral Asgard *thyA*; a process known as “non-homologous displacement”^29^. The non-homologous displacement of the *thyA* gene by *thyX* has already been suggested for diverse bacterial and archaeal species^2, 10^, but genetic (Fig. 5) and biochemical (Fig. 6) validation of the functionality of the acquired gene via an interkingdom LGT has been lacking.

The phylogeny of *Psyn* ThyX, together with its biochemical properties, prompted us to investigate more globally the distribution and phylogeny of folate-dependent enzymes in Asgard archaea. We first observed that *Psyn* carries many folate-dependent enzymes, which raised the possibility that a complex interdependent reaction network operates in this species to chemically activate and transfer one-carbon units. The detailed reconstruction of the *Psyn* one-carbon metabolism (Fig. 8) is fully consistent with the reported lifestyle of this species in anaerobic marine methane seep sediments. In particular, a syntrophic amino acid utilization of *Psyn* using serine, glycine and histidine is highly feasible, as these amino acids can readily provide one carbon units for the folate metabolism at different oxidation levels. *Psyn* likely uses folate derivatives for the synthesis of thymidylate, purine, and methionine. In many cases, a strong association of the genes for the folate-dependent enzymes with additional functionally linked genes, for instance, glycine cleavage or histidine catabolism, was also observed (Fig. 8b). We also noticed that, in agreement with a previous study^18^, biosynthetic pathways for folic acid and tetrahydromethanopterin appear incomplete in *Psyn*. However, the growth medium of *Psyn* is supplemented with folic acid-containing vitamin supplement^18^, and many bacteria are known to transport folates^30, 31, 32^. These observations raised the possibility of *Psyn* cross-feeding with the folates produced by its symbiotic partners, as has been previously observed for bacterial communities^31, 33^.

Strikingly, most, if not all, folate-dependent *Psyn* enzymes appear to have a bacterial origin (Figs. 4 and 7). The transfer of genes from Bacteria to Archaea is well documented^34^. However, our study has revealed a remarkable multigene transfer that globally influences the central metabolism of this enigmatic species. Our phylogenetic analyses did not suggest a well-defined single donor for these transferred genes that are scattered around the *Psyn* genome. Therefore, these transfers likely have occurred on multiple and sequential occasions. Our observations agree well with the previous suggestion that LGT of bacterial metabolic networks may provide an adaptive benefit to *Psyn* in response to changing environments^35^. Our observations are of general interest as our HGTector analysis (Fig. 3) indicated that at least 4% of *Psyn* genes correspond to transfer events including many genes implicated *e*.*g*. in amino acid and nucleotide metabolism. Our analyses may underestimate the true extent of LGT for this species, as the initial discovery of the first Asgard genome -a lokiarchaeal species-reported a substantial fraction of genes with bacterial affinities (29% of all genes)^36^. Widespread interdomain LGTs have also been proposed in uncultured planktonic thaumarchaeota and euryarchaeota^37^. Association of *Psyn* with co-cultured bacteria might have favoured the gene exchanges, possibly influencing microbial population structure under natural conditions^38^. Several different archaeal molecular mechanisms facilitating DNA transfer have been described^39^, providing plausible means for LGT between Bacteria and Asgard archaea. Finally, frequent LGTs between *Psyn* and different bacterial species might partially explain the apparent chimeric profiles of Asgard genomes^40^, which seemingly result from the frequent mixing of ancestral archaeal genes with laterally inherited bacterial ones.

Regarding the eukaryogenesis process, the possible contribution of Asgard to the eukaryotic folate metabolism remains elusive. Indeed, our results support the existence of multiple and independent lateral gene transfers from diverse bacteria to Asgard, blurring an exact scenario. Nevertheless, the ThyA family of thymidylate synthases appears ancestral in Asgard considering the phylogenetic positioning of the Asgard sequences deeply nested in the bacterial phylogenetic tree. Our analyses also indicate that eukaryotic ThyA does not derive from Asgard archaea but also has a bacterial origin. Both scenarios suggest that the present-day thymidylate metabolism in Eukarya and Asgard archaea originated in bacteria. It was recently proposed that the contribution of Asgard to the present eukaryotic gene repertoire is very limited^41^. However, widespread recent LGTs events from bacteria to Asgard displacing the original archaeal genes would underestimate the contribution of the current day Asgard to the eukaryotic gene repertoire. In turn, our study on the thymidylate metabolism might represent only the tip of an iceberg of functional bacterial genes in Asgard genomes.

Taken together, our data support the idea that Asgard archaea harbor highly mosaic genomes that have received many bacterial genes, even for central metabolic processes such as the folate metabolism. We have also experimentally demonstrated that some of the transferred *Psyn* genes, like *Psyn thyX*, are fully functional *in vivo* and *in vitro*. More general studies on Asgard gene evolution are now needed to better appreciate the pervasiveness of interdomain LGT, as well as the origin of the eukaryotic gene repertoire in this fascinating group of prokaryotes.

## Materials and Methods

### Bioinformatics methods and genome-wide analyses

The complete genome sequence available for the *Candidatus* Prometheoarchaeum syntrophicum strain MK-D1 (*Psyn*) chromosome (NZ_CP042905.1) was used for bioinformatics analyses. This genome is 4,278 Mb with a GC content of 31.2%. RPKM values for gene expression were determined using the Geneious Prime® 2021.1.1 (Build 2021-03-12). Metabolic reconstructions were performed using the RAST server and KEGG databases. These analyses were complemented with highly sensitive manual similarity searches using HHpred^42^.

### Structural alignments and structure modelling

<u>PRO</u>file <u>M</u>ultiple <u>A</u>lignment with predicted <u>L</u>ocal <u>S</u>tructures and <u>3D</u> constraints (PROMALS3D) was used to align the *Psyn* ThyX sequence with the *Thermotoga (T*.*) maritima* ThyX structure (PDB 5CHP). PROMALS3D (http://prodata.swmed.edu/promals3d/promals3d.php) aligns multiple protein sequences and/or structures, with enhanced information from database searches, secondary structure prediction, and 3D structures. Input sequences and structures used FASTA and PDB formats, respectively. (red: alpha-helix, blue: beta-strand).

A model of the *Psyn* ThyX homotetramer was constructed with the protein structure modeling program Modeller using PDB structures 1O26, 3N0B, and 6J61 as templates^21^. Ten initial models were constructed and the best one was chosen using molpdf and DOPE score, and by evaluating the stereochemical quality with PROCHECK. A structural superposition of the model with PDB structure 3GT9 using the Chimera software^22^ allowed the addition of FAD, dUMP and folate molecules. To improve the structural model obtained, molecular dynamics simulations were performed with CHARMM^43^ and Namd^44^ molecular mechanics softwares.

### Automated prediction of LGT

Automated analyses of potential LGT events were performed using a HGTector^23^. This method first performs Diamond all-against-all similarity searches of each protein-coding sequence of *Psyn*. The NCBI genome database was downloaded and compiled (on August 2021) on a local machine. The output of the similarity searches was passed to the analysis program HGTector where the taxonomy of the self-group was defined as *Candidatus* Prometheoarchaeum syntrophicum (NCBI TaxID: 2594042). The close-group was automatically inferred by HGTector as the “superkingdom Archaea” (TaxID: 2157) that contains 1062 taxa in the reference database. From the statistical distribution of close and distal hits, HGTector predicted a list of potentially transferred genes. The output of the program is a scatter plot, where the horizontal axis represents the close score, the vertical axis represents the distal score, and potential LGT genes are colored in yellow. We slightly modified the source code which was made publicly available at: https://github.com/cgneo/neoHGT.

### Phylogenetic analyses

For phylogenetic analyses, we constructed a database with a backbone of 183 *thyX* sequences that cover all the major bacterial and archaeal phyla derived from a previous study^10^. These were submitted to BLASTP, with an *e*-value cut-off of 1e-05, against the *Psyn* genome, as well as against the 140 metagenomic bins larger than 2Mb assigned to Asgard archaea publicly available in the Genbank database. We used similar approaches with the other folate enzymes with 355 ThyA, 53 FTHFS, 40 MetH, 58 PurH, 70 FolA, 85 FolD, 33 MTHFR and 49 MTHFS sequences to find homologous sequences in Asgard genomes. Homologs found in Asgard genomes were then used as seeds for reciprocal BLAST searches against an NR database (15/01/2021 version) to find the best homolog outside Asgards. Identical sequences (same identifying code or 100% sequence similarity) were finally removed to generate the final dataset.

For analysis of the genomic environment of the *Psyn thyX* gene, we collected all the genes located 15 kb downstream and 15 kb upstream of the *Psyn thyX* genes and searched for homologs with BLASTP against an NR database (15/01/2021 version). The first BLASTP hit was retrieved, as well as the percentages of the archaeal sequences in the taxonomy profile of the first top 100 hits.

Phylogenetic analyses were performed using the obtained protein sequence data sets (see above) that were aligned using MAFFT v7.388 with default settings^45^. Identical sequences belonging to the same phylum or genus were removed to retain a single representative sequence. Ambiguously aligned sites were removed using trimAl v1.4.rev15^46^ with the “-automated1” and “-phylip” options. The phylogenetic position of the Asgard sequences was then inferred with PhyML v. 1.8.1^47^ using the best substitution models as determined by Smart Model Selection^48^, and 1000 bootstrapped data sets were used to estimate the statistical confidences of the nodes. Trees were then visualized using the FigTree software (https://github.com/rambaut/figtree/releases). Raw data including BLAST outputs, table with the accession numbers of each Asgard sequences, alignment files, alignments used for the phylogenies, Newick tree files and additional information can be downloaded at http://gofile.me/2ppPR/HvmT893hn.

### Design and synthesis of *Psyn* ThyX expression plasmid

pTwist-Psyn-ThyX plasmid (3370bp), carrying the synthetic *Psyn thyX* gene of 897bp was synthesized by Twist Biosciences (https://www.twistbioscience.com) and confirmed by sequencing. In the final design, the target gene *Psyn thyX* was placed under the control of a strong bacteriophage T5 promotor carrying *lacO* sites. This construct also carried an appropriate ribosome binding site, a lambda t0 transcriptional terminator, and the Amp^R^ gene. The pTwist-Psyn-ThyX plasmid uses a p15A origin of replication and contains the codon-optimized synthetic gene of *Psyn*-ThyX with an N-terminal 6xHis sequence.

### Genetic complementation tests

The ability of *Psyn thyX* to permit thymidine-independent growth of the *E. coli* thymidine-auxotroph FE013 strain (Δ*thyA::aphA3 lacI*, derived from wild type MG1655 strain)^10^ was scored after three days at room temperature or at 37°C in the presence of 1mM IPTG using either minimal M9 (shown in Figure 5) or thymidine-deprived enriched medium L^+^. Eight individual colonies all demonstrating similar phenotypes in the absence of thymidine were tested.

### Protein purification

*E. coli* FE013 cells carrying pTwist-Psyn-ThyX were grown at 37°C and shaken at 150rpm in liquid Luria Bertani medium supplemented with ampicillin (final 100 μg/mL) until an absorbance of 600 nm around 0.7 was reached. Expression of recombinant *Psyn* ThyX was induced by the addition of 0.5 mM IPTG (isopropyl-b-D-thiogalactopyranoside) during 3 hours at 37°C. Cells were harvested by centrifugation at 6000 x g at 4°C for 30min before storage at -20°C. The *Psyn* ThyX recombinant protein was purified on Protino Ni-TED column (Macherey-Nagel) as previously described^49^. Stepwise protein elution was performed with imidazole at 50mM, 100mM, 150mM and 250mM in a phosphate buffer (50mM Na_2_HPO_4_-NaH_2_PO_4_, pH8, 300mM NaCl). The 100mM imidazole fractions containing the *Psyn* ThyX enzyme were pooled, buffer exchanged on Econo-Pac PD-10 Columns (Bio-Rad Laboratories, Hercules, CA) with phosphate buffer (50mM Na_2_HPO_4_-NaH_2_PO_4_, pH8, 300mM NaCl), concentrated to a final concentration of 4μM and stored at -20°C in the same phosphate buffer complemented with glycerol (10% v/v).

The purified proteins were analyzed on 12% TGX Precast Gels (Bio-Rad Laboratories, Hercules, CA) followed by Coomassie Brillant Blue staining. For immunoblotting detection, after protein transfer, the nitrocellulose membranes (Bio-Rad) were treated with 20% blocking buffer (Li-Cor Biosciences, Lincoln, US) followed by incubation with anti-His-tag mouse primary antibody (dilution1/5000, Bio-Rad) and revelation with a IRDye 800CW Goat anti-mouse IgG secondary antibody (dilution 1/10000, Li-Cor Biosciences) using a ChemiDoc− Touch Imaging System (Bio-Rad).

### Enzymatic activity tests

The NADPH oxidase assay consists in measuring the conversion of NADPH to NADP^+^ via the decrease in absorbance at 340 nm. During the test, in 96-well plates, one hundred microlitres of the standard reaction mixture (50mM Buffer Na_2_HPO_4_-NaH_2_PO_4_ pH 8, NaCl 300 mM, MgCl_2_ 2 mM, FAD 50 μM, β-mercaptoethanol 1.43 mM, dUMP 20 μM, NADPH 750 μM, glycerol 8%, 0.4 μM of purified *Psyn*ThyX) were incubated in a Chameleon II microplate reader (Hidex) at 25°C. Measurements took place over 15 minutes and were done in triplicates. The enzyme-free reaction was used as a negative control.

Tests of ThyX inhibition were performed by incubation of molecules at 100 or 200μM with *Psyn*ThyX enzyme (0.4 μM) in the standard reaction mixture, without NADPH, for 10 min at 25°C before starting measurements by automatically injecting NADPH. The molecules “2716” and C8C1 were solubilized in dimethylsulfoxide (DMSO) and used at 1% final concentration of DMSO during the test. % of inhibition was calculated using the following equation: ((Vo-Vi)/Vo)*100; Vo and Vi are, respectively, the initial rates of the reaction without or with addition of molecule to the assay.

## Acknowledgements

We thank CNRS, INSERM, and E. Polytechnique for their financial support. X.L is a student of the Bachelor of Science program of E. Polytechnique.

## Author contributions

J.F., H.B., U.L, and H.M. conceived the study and wrote the manuscript with contributions from all other authors. J.F. and H.M performed phylogenetic and bioinformatics analyses. H.B and L.M provided biochemical data and U.L and H.M performed genetic experiments. J-C.L. provided a structural model and Z.L performed HGTector analyses. All authors read and commented on the manuscript.

## Additional information

Supplementary information for this manuscript includes the supplementary figures 1-9 and the supplementary table 1.

## References

1. Iyer LM, et al. Quod erat demonstrandum? The mystery of experimental validation of apparently erroneous computational analyses of protein sequences. Genome Biol 2, RESEARCH0051 (2001).

2. Myllykallio H, Lipowski G, Leduc D, Filee J, Forterre P, Liebl U. An alternative flavin-dependent mechanism for thymidylate synthesis. Science 297, 105–107 (2002).

3. Giladi M, Bitan-Banin G, Mevarech M, Ortenberg R. Genetic evidence for a novel thymidylate synthase in the halophilic archaeon Halobacterium salinarum and in Campylobacter jejuni. FEMS Microbiol Lett 216, 105–109 (2002).

4. Myllykallio H, Leduc D, Filee J, Liebl U. Life without dihydrofolate reductase FolA. Trends Microbiol 11, 220–223 (2003).

5. Leduc D, et al. Functional evidence for active site location of tetrameric thymidylate synthase X at the interphase of three monomers. Proc Natl Acad Sci U S A 101, 7252–7257 (2004).

6. Carreras CW, Santi DV. The catalytic mechanism and structure of thymidylate synthase. Annu Rev Biochem 64, 721–762 (1995).

7. Schober AF, et al. A Two-Enzyme Adaptive Unit within Bacterial Folate Metabolism. Cell Rep 27, 3359–3370 e3357 (2019).

8. Escartin F, Skouloubris S, Liebl U, Myllykallio H. Flavin-dependent thymidylate synthase X limits chromosomal DNA replication. Proc Natl Acad Sci U S A 105, 9948–9952 (2008).

9. Myllykallio H, Sournia P, Heliou A, Liebl U. Unique Features and Anti-microbial Targeting of Folate- and Flavin-Dependent Methyltransferases Required for Accurate Maintenance of Genetic Information. Front Microbiol 9, 918 (2018).

10. Stern A, Mayrose I, Penn O, Shaul S, Gophna U, Pupko T. An evolutionary analysis of lateral gene transfer in thymidylate synthase enzymes. Syst Biol 59, 212–225 (2010).

11. Giovannoni SJ, et al. Genome streamlining in a cosmopolitan oceanic bacterium. Science 309, 1242–1245 (2005).

12. Arnoriaga-Rodriguez M, et al. Obesity-associated deficits in inhibitory control are phenocopied to mice through gut microbiota changes in one-carbon and aromatic amino acids metabolic pathways. Gut, (2021).

13. Krone UE, McFarlan SC, Hogenkamp HP. Purification and partial characterization of a putative thymidylate synthase from Methanobacterium thermoautotrophicum. Eur J Biochem 220, 789–794 (1994).

14. Nyce GW, White RH. dTMP biosynthesis in Archaea. J Bacteriol 178, 914–916 (1996).

15. Maden BE. Tetrahydrofolate and tetrahydromethanopterin compared: functionally distinct carriers in C1 metabolism. Biochem J 350 Pt 3, 609–629 (2000).

16. Adam PS, Borrel G, Gribaldo S. An archaeal origin of the Wood-Ljungdahl H4MPT branch and the emergence of bacterial methylotrophy. Nat Microbiol 4, 2155–2163 (2019).

17. Zaremba-Niedzwiedzka K, et al. Asgard archaea illuminate the origin of eukaryotic cellular complexity. Nature 541, 353–358 (2017).

18. Imachi H, et al. Isolation of an archaeon at the prokaryote-eukaryote interface. Nature 577, 519–525 (2020).

19. Liu Y, et al. Expanded diversity of Asgard archaea and their relationships with eukaryotes. Nature 593, 553–557 (2021).

20. Williams TA, Cox CJ, Foster PG, Szollosi GJ, Embley TM. Phylogenomics provides robust support for a two-domains tree of life. Nat Ecol Evol 4, 138–147 (2020).

21. Webb B, Sali A. Comparative Protein Structure Modeling Using MODELLER. Curr Protoc Protein Sci 86, 2 9 1-2 9 37 (2016).

22. Pettersen EF, et al. UCSF Chimera--a visualization system for exploratory research and analysis. J Comput Chem 25, 1605–1612 (2004).

23. Zhu Q, Kosoy M, Dittmar K. HGTector: an automated method facilitating genome-wide discovery of putative horizontal gene transfers. BMC Genomics 15, 717 (2014).

24. Kanehisa M, Sato Y, Morishima K. BlastKOALA and GhostKOALA: KEGG Tools for Functional Characterization of Genome and Metagenome Sequences. J Mol Biol 428, 726–731 (2016).

25. Graziani S, et al. Functional analysis of FAD-dependent thymidylate synthase ThyX from Paramecium bursaria Chlorella virus-1. J Biol Chem 279, 54340–54347 (2004).

26. Graziani S, et al. Catalytic mechanism and structure of viral flavin-dependent thymidylate synthase ThyX. J Biol Chem 281, 24048–24057 (2006).

27. Abu El Asrar R, et al. Discovery of a new Mycobacterium tuberculosis thymidylate synthase X inhibitor with a unique inhibition profile. Biochem Pharmacol 135, 69–78 (2017).

28. Modranka J, et al. Synthesis and Structure-Activity Relationship Studies of Benzo[b][1,4]oxazin-3(4H)-one Analogues as Inhibitors of Mycobacterial Thymidylate Synthase X. ChemMedChem 14, 645–662 (2019).

29. Galperin MY, Koonin EV. Sources of systematic error in functional annotation of genomes: domain rearrangement, non-orthologous gene displacement and operon disruption. In Silico Biol 1, 55–67 (1998).

30. Shane B, Stokstad EL. Transport and metabolism of folates by bacteria. J Biol Chem 250, 2243–2253 (1975).

31. Graber JR, Breznak JA. Folate cross-feeding supports symbiotic homoacetogenic spirochetes. Appl Environ Microbiol 71, 1883–1889 (2005).

32. Leduc D, et al. Flavin-dependent thymidylate synthase ThyX activity: implications for the folate cycle in bacteria. J Bacteriol 189, 8537–8545 (2007).

33. Soto-Martin EC, et al. Vitamin Biosynthesis by Human Gut Butyrate-Producing Bacteria and Cross-Feeding in Synthetic Microbial Communities. mBio 11, (2020).

34. Gogarten JP, Doolittle WF, Lawrence JG. Prokaryotic evolution in light of gene transfer. Mol Biol Evol 19, 2226–2238 (2002).

35. Pal C, Papp B, Lercher MJ. Adaptive evolution of bacterial metabolic networks by horizontal gene transfer. Nat Genet 37, 1372–1375 (2005).

36. Spang A, et al. Complex archaea that bridge the gap between prokaryotes and eukaryotes. Nature 521, 173–179 (2015).

37. Deschamps P, Zivanovic Y, Moreira D, Rodriguez-Valera F, Lopez-Garcia P. Pangenome evidence for extensive interdomain horizontal transfer affecting lineage core and shell genes in uncultured planktonic thaumarchaeota and euryarchaeota. Genome Biol Evol 6, 1549–1563 (2014).

38. Polz MF, Alm EJ, Hanage WP. Horizontal gene transfer and the evolution of bacterial and archaeal population structure. Trends Genet 29, 170–175 (2013).

39. Wagner A, et al. Mechanisms of gene flow in archaea. Nat Rev Microbiol 15, 492–501 (2017).

40. Garg SG, et al. Anomalous Phylogenetic Behavior of Ribosomal Proteins in Metagenome-Assembled Asgard Archaea. Genome Biol Evol 13, (2021).

41. Knopp M, Stockhorst S, van der Giezen M, Garg SG, Gould SB. The Asgard Archaeal-Unique Contribution to Protein Families of the Eukaryotic Common Ancestor Was 0.3. Genome Biol Evol 13, (2021).

42. Zimmermann L, et al. A Completely Reimplemented MPI Bioinformatics Toolkit with a New HHpred Server at its Core. J Mol Biol 430, 2237–2243 (2018).

43. Brooks BR, et al. CHARMM: the biomolecular simulation program. J Comput Chem 30, 1545–1614 (2009).

44. Phillips JC, et al. Scalable molecular dynamics with NAMD. J Comput Chem 26, 1781–1802 (2005).

45. Katoh K, Standley DM. MAFFT multiple sequence alignment software version 7: improvements in performance and usability. Mol Biol Evol 30, 772–780 (2013).

46. Capella-Gutierrez S, Silla-Martinez JM, Gabaldon T. trimAl: a tool for automated alignment trimming in large-scale phylogenetic analyses. Bioinformatics 25, 1972–1973 (2009).

47. Guindon S, Dufayard JF, Lefort V, Anisimova M, Hordijk W, Gascuel O. New algorithms and methods to estimate maximum-likelihood phylogenies: assessing the performance of PhyML 3.0. Syst Biol 59, 307–321 (2010).

48. Lefort V, Longueville JE, Gascuel O. SMS: Smart Model Selection in PhyML. Mol Biol Evol 34, 2422–2424 (2017).

49. Ulmer JE, Boum Y, Thouvenel CD, Myllykallio H, Sibley CH. Functional analysis of the Mycobacterium tuberculosis FAD-dependent thymidylate synthase, ThyX, reveals new amino acid residues contributing to an extended ThyX motif. J Bacteriol 190, 2056–2064 (2008).

