## Supplementary figures for "Bacterial origin of thymidylate and folate metabolism in Asgard Archaea"

### Slide 1
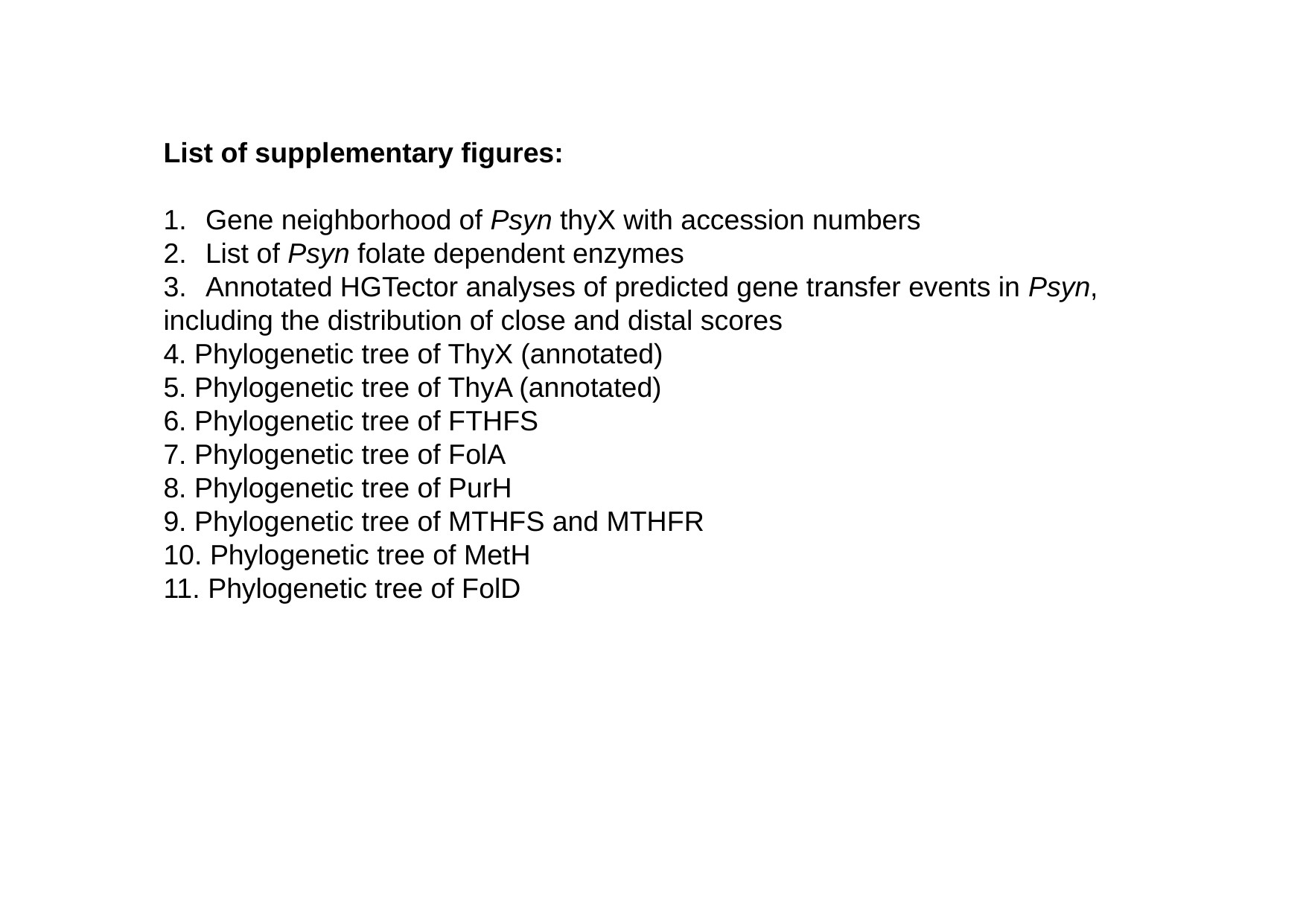

List of supplementary figures:
Gene neighborhood of Psyn thyX with accession numbers
List of Psyn folate dependent enzymes
Annotated HGTector analyses of predicted gene transfer events in Psyn,
including the distribution of close and distal scores
4. Phylogenetic tree of ThyX (annotated)
5. Phylogenetic tree of ThyA (annotated)
6. Phylogenetic tree of FTHFS
7. Phylogenetic tree of FolA
8. Phylogenetic tree of PurH
9. Phylogenetic tree of MTHFS and MTHFR
10. Phylogenetic tree of MetH
11. Phylogenetic tree of FolD

### Slide 2
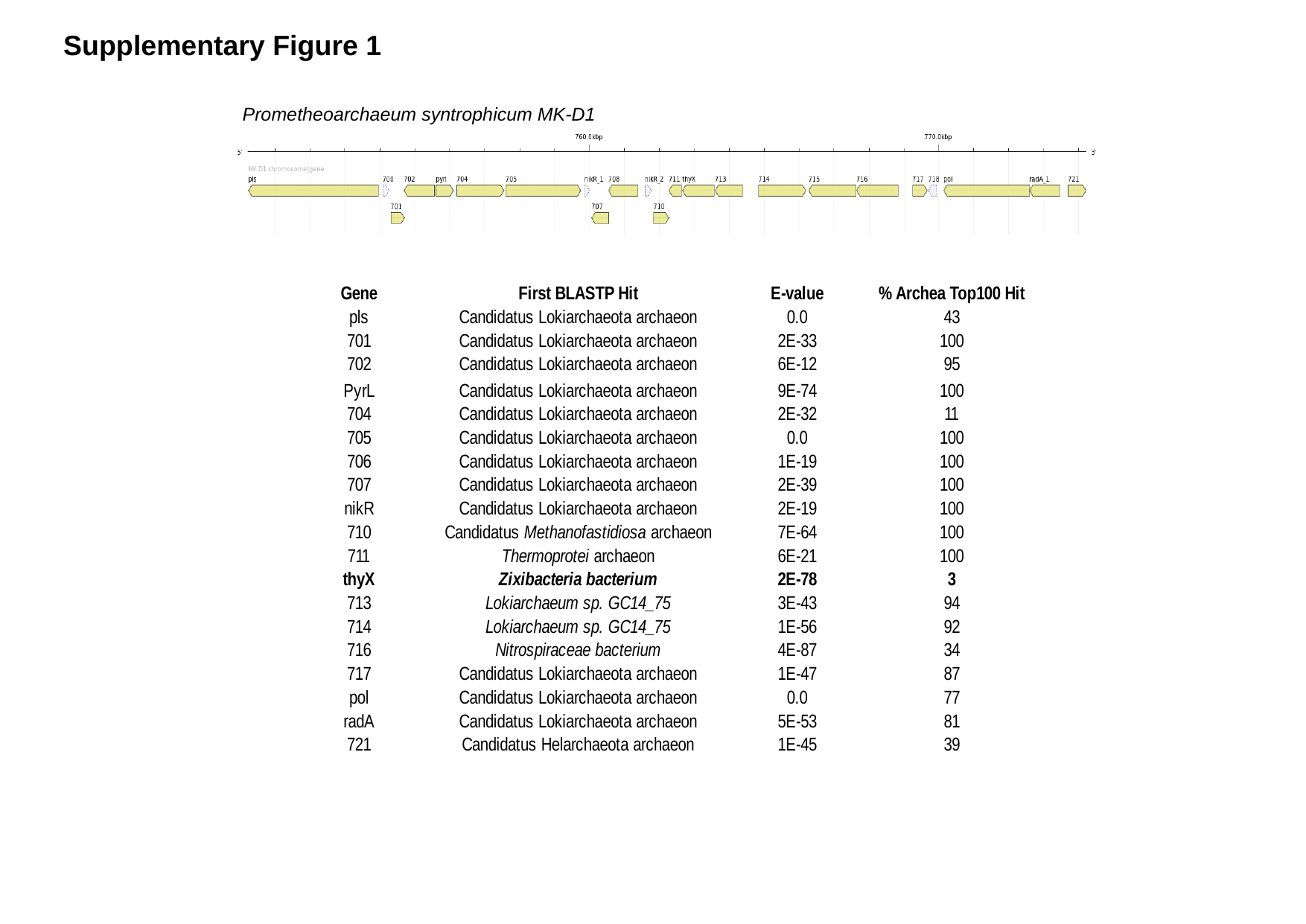

Supplementary Figure 1
Prometheoarchaeum syntrophicum MK-D1

### Slide 3
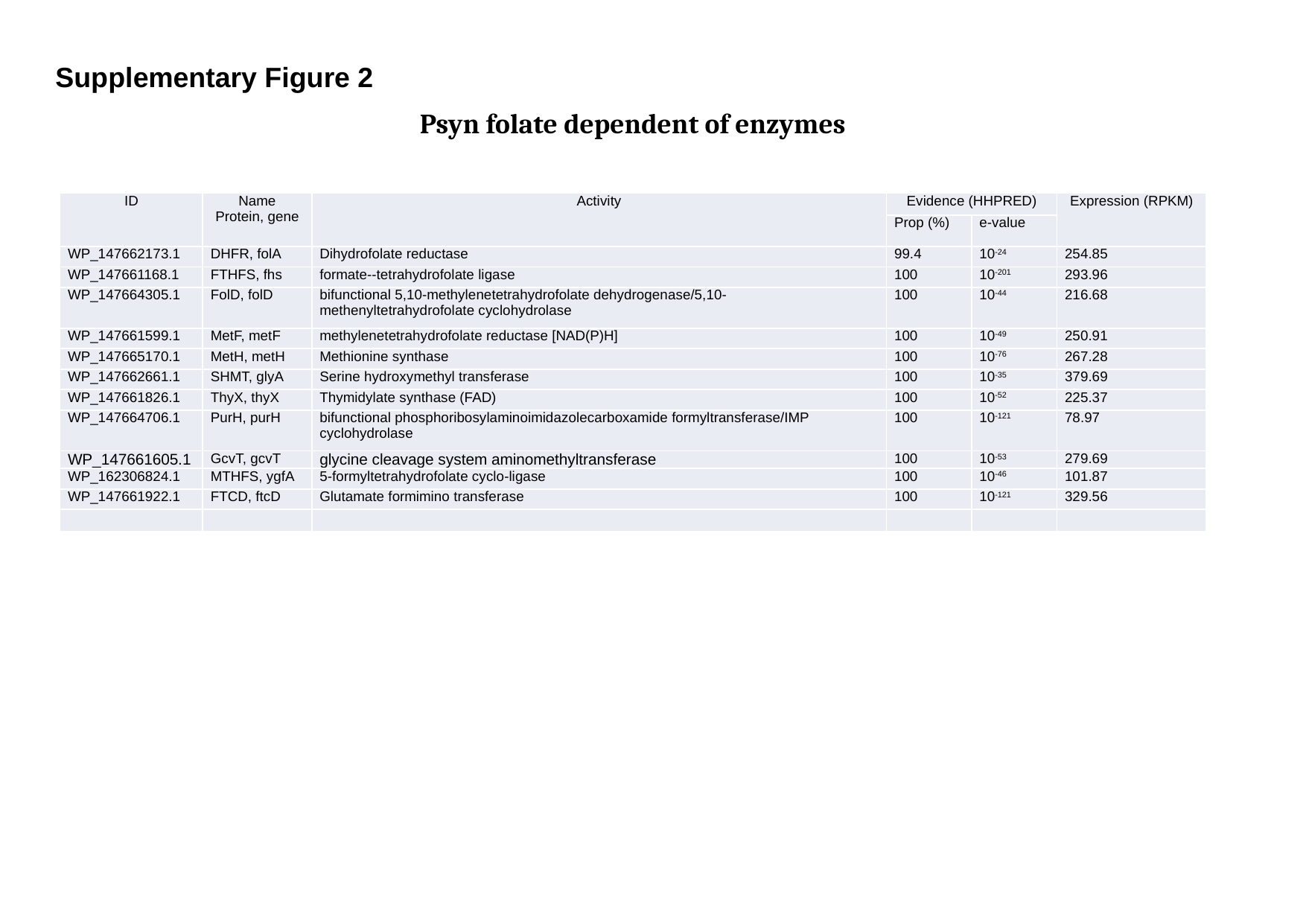

Supplementary Figure 2
Psyn folate dependent of enzymes
| ID | Name Protein, gene | Activity | Evidence (HHPRED) | | Expression (RPKM) |
| --- | --- | --- | --- | --- | --- |
| | | | Prop (%) | e-value | |
| WP\_147662173.1 | DHFR, folA | Dihydrofolate reductase | 99.4 | 10^-24^ | 254.85 |
| WP\_147661168.1 | FTHFS, fhs | formate--tetrahydrofolate ligase | 100 | 10^-201^ | 293.96 |
| WP\_147664305.1 | FolD, folD | bifunctional 5,10-methylenetetrahydrofolate dehydrogenase/5,10-methenyltetrahydrofolate cyclohydrolase | 100 | 10^-44^ | 216.68 |
| WP\_147661599.1 | MetF, metF | methylenetetrahydrofolate reductase [NAD(P)H] | 100 | 10^-49^ | 250.91 |
| WP\_147665170.1 | MetH, metH | Methionine synthase | 100 | 10^-76^ | 267.28 |
| WP\_147662661.1 | SHMT, glyA | Serine hydroxymethyl transferase | 100 | 10^-35^ | 379.69 |
| WP\_147661826.1 | ThyX, thyX | Thymidylate synthase (FAD) | 100 | 10^-52^ | 225.37 |
| WP\_147664706.1 | PurH, purH | bifunctional phosphoribosylaminoimidazolecarboxamide formyltransferase/IMP cyclohydrolase | 100 | 10^-121^ | 78.97 |
| WP\_147661605.1 | GcvT, gcvT | glycine cleavage system aminomethyltransferase | 100 | 10^-53^ | 279.69 |
| WP\_162306824.1 | MTHFS, ygfA | 5-formyltetrahydrofolate cyclo-ligase | 100 | 10^-46^ | 101.87 |
| WP\_147661922.1 | FTCD, ftcD | Glutamate formimino transferase | 100 | 10^-121^ | 329.56 |

### Slide 4
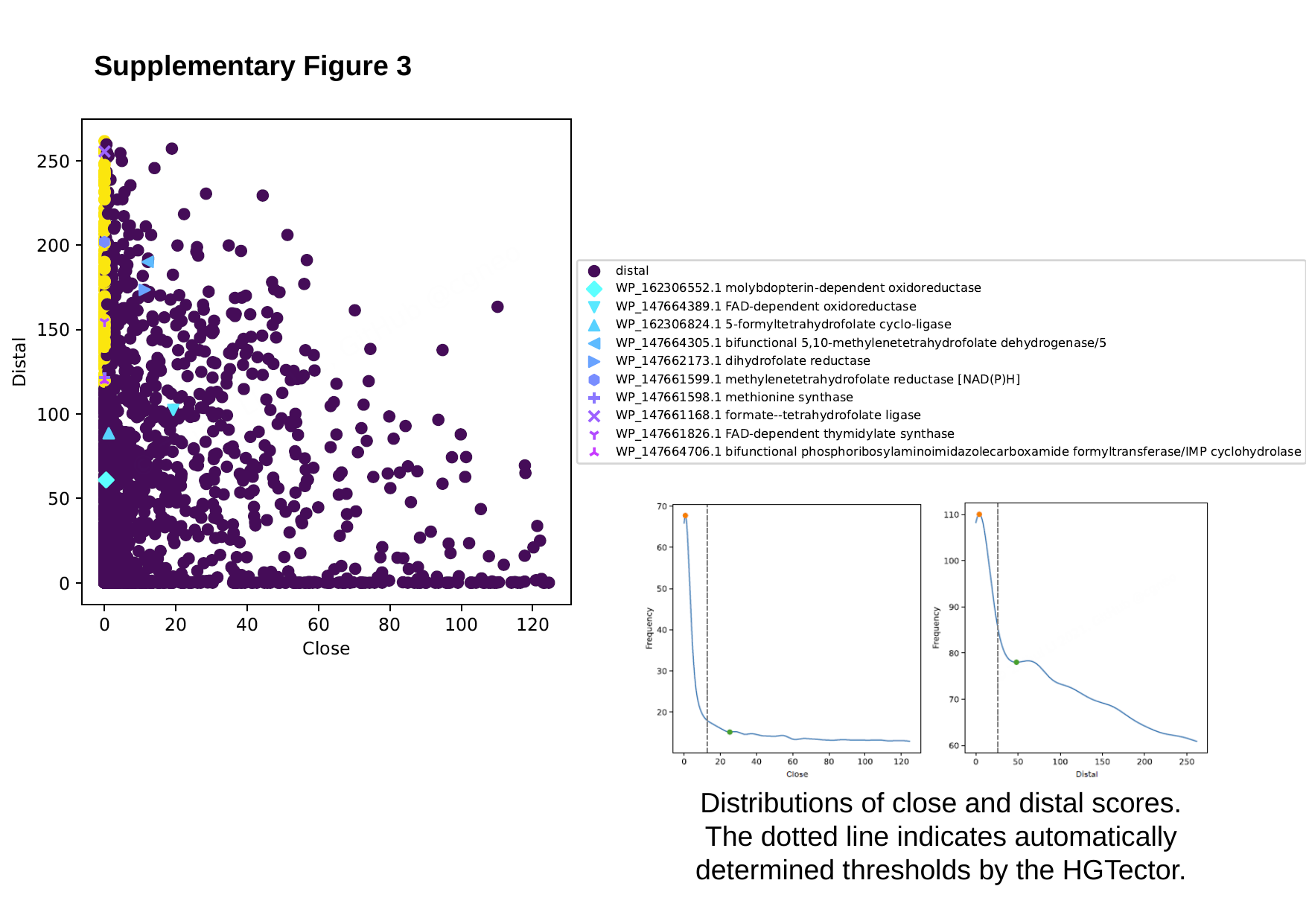

Supplementary Figure 3
Distributions of close and distal scores. The dotted line indicates automatically determined thresholds by the HGTector.

### Slide 5
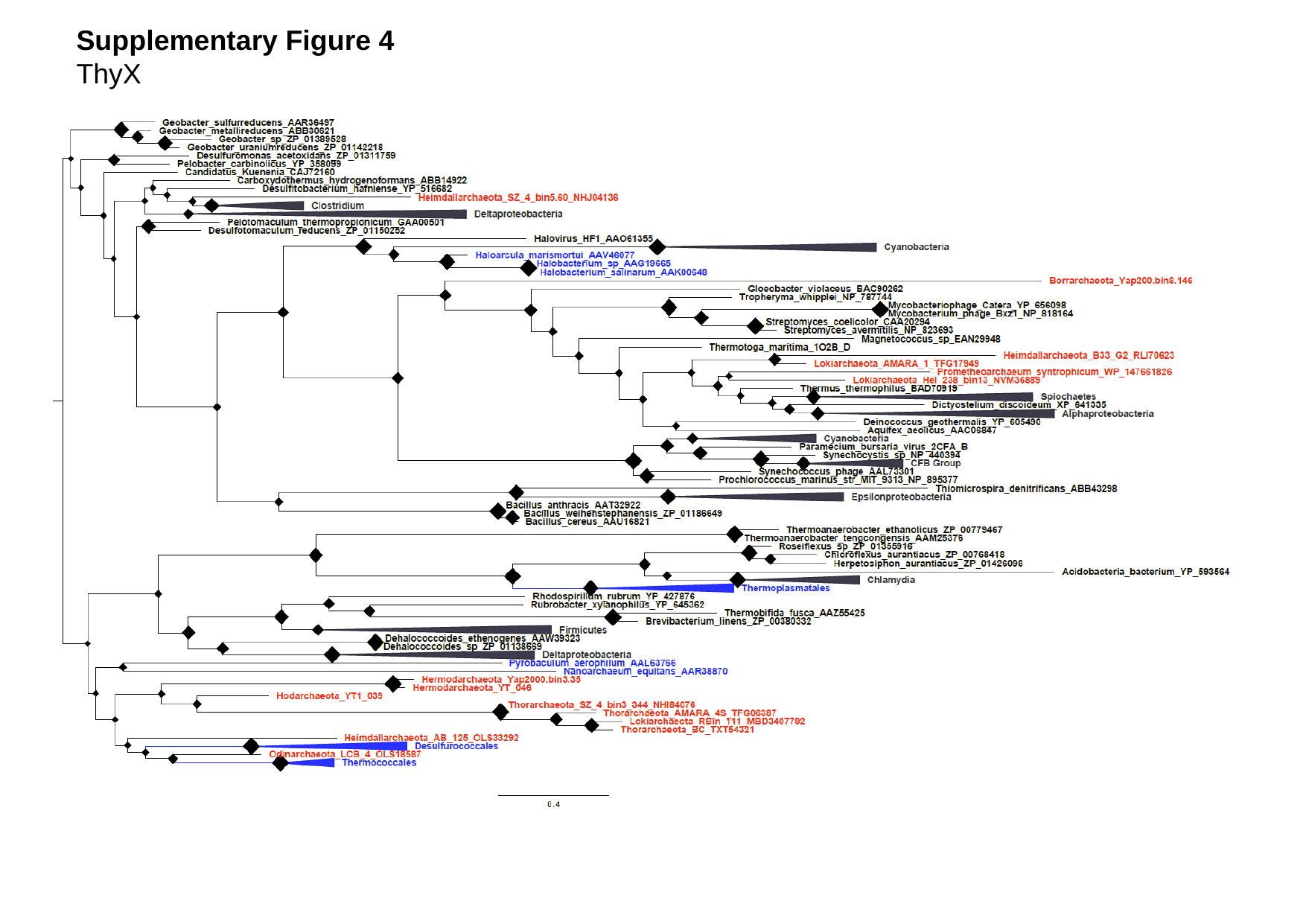

Supplementary Figure 4
ThyX

### Slide 6
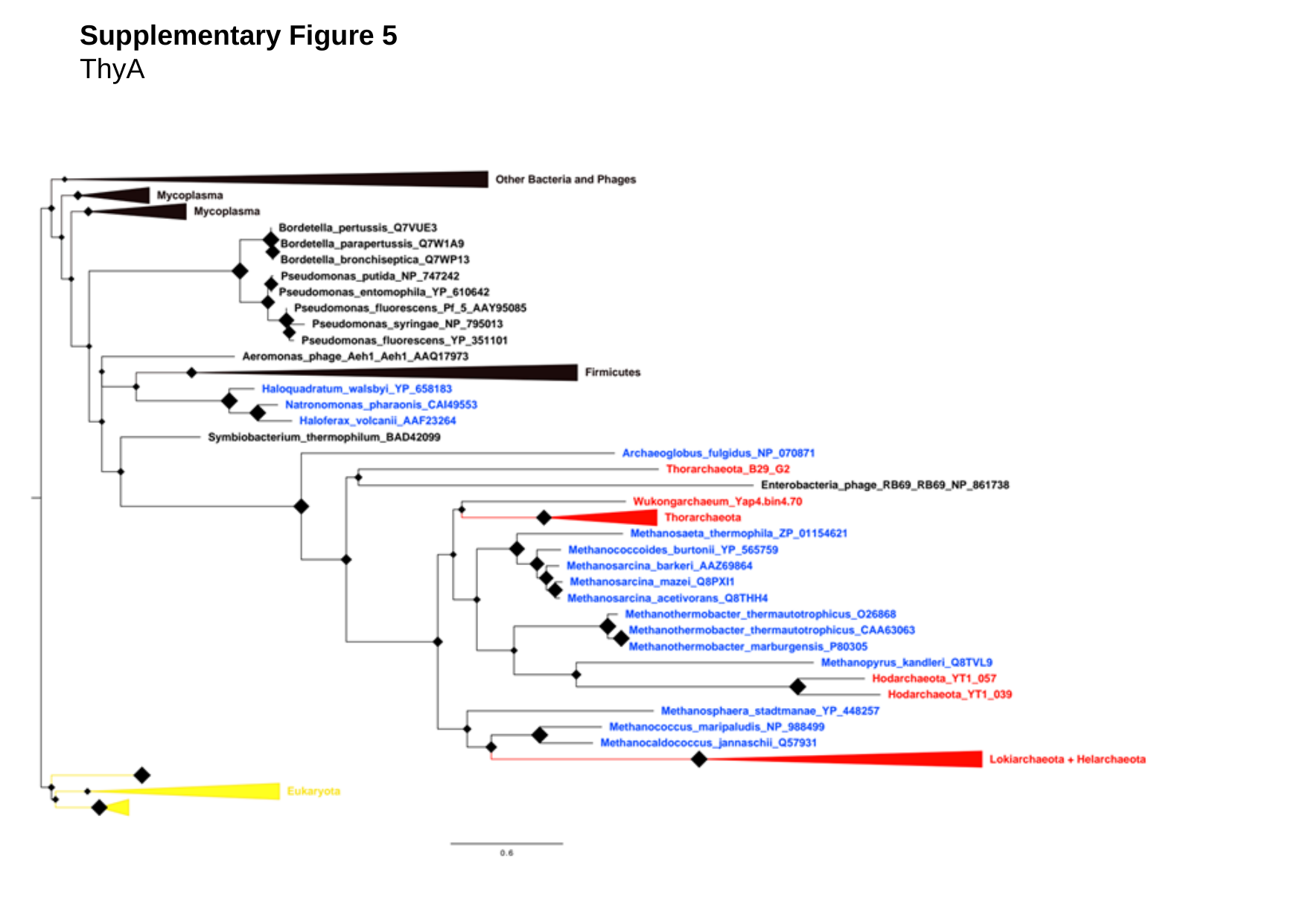

Supplementary Figure 5
ThyA

### Slide 7
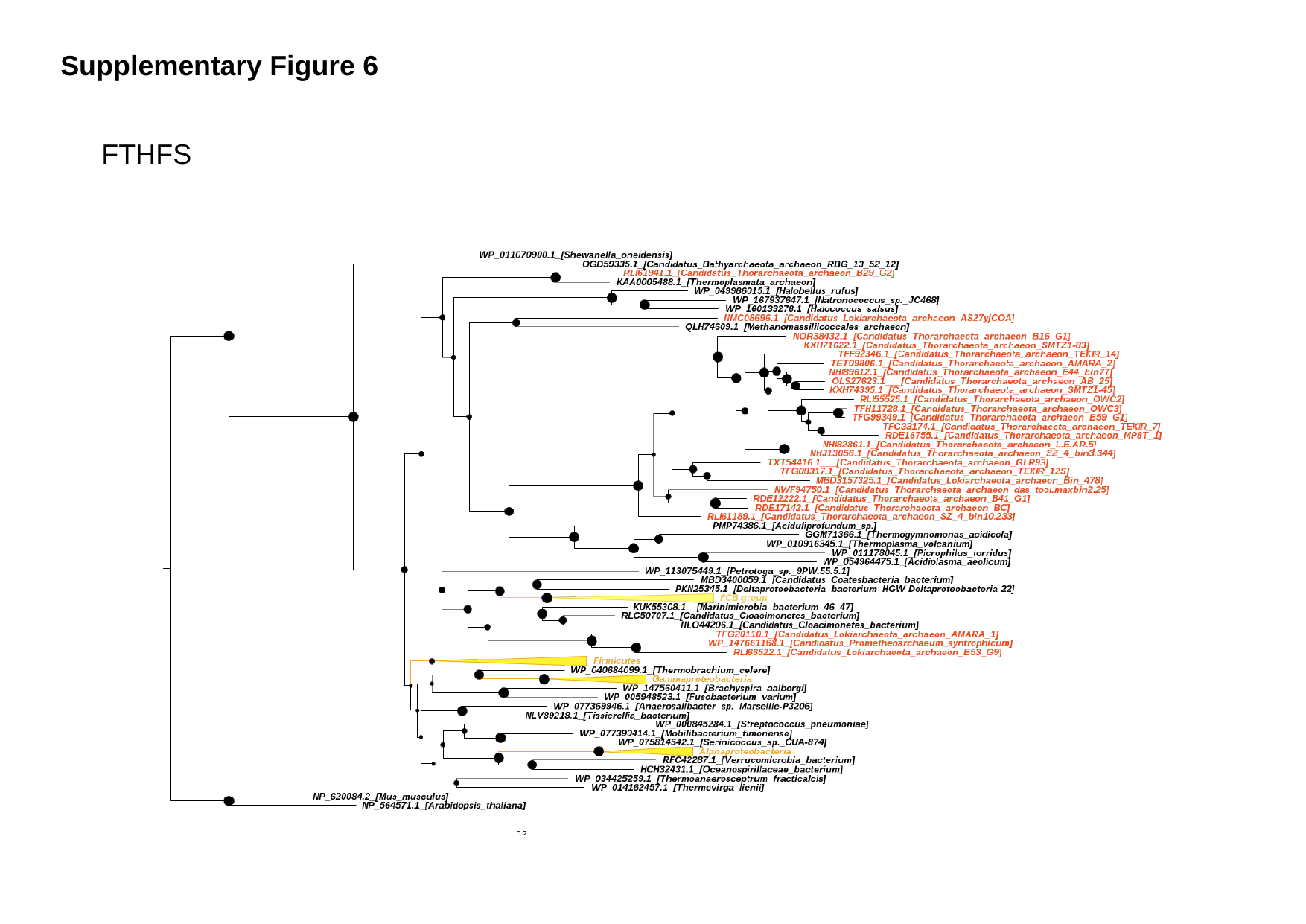

Supplementary Figure 6
FTHFS

### Slide 8
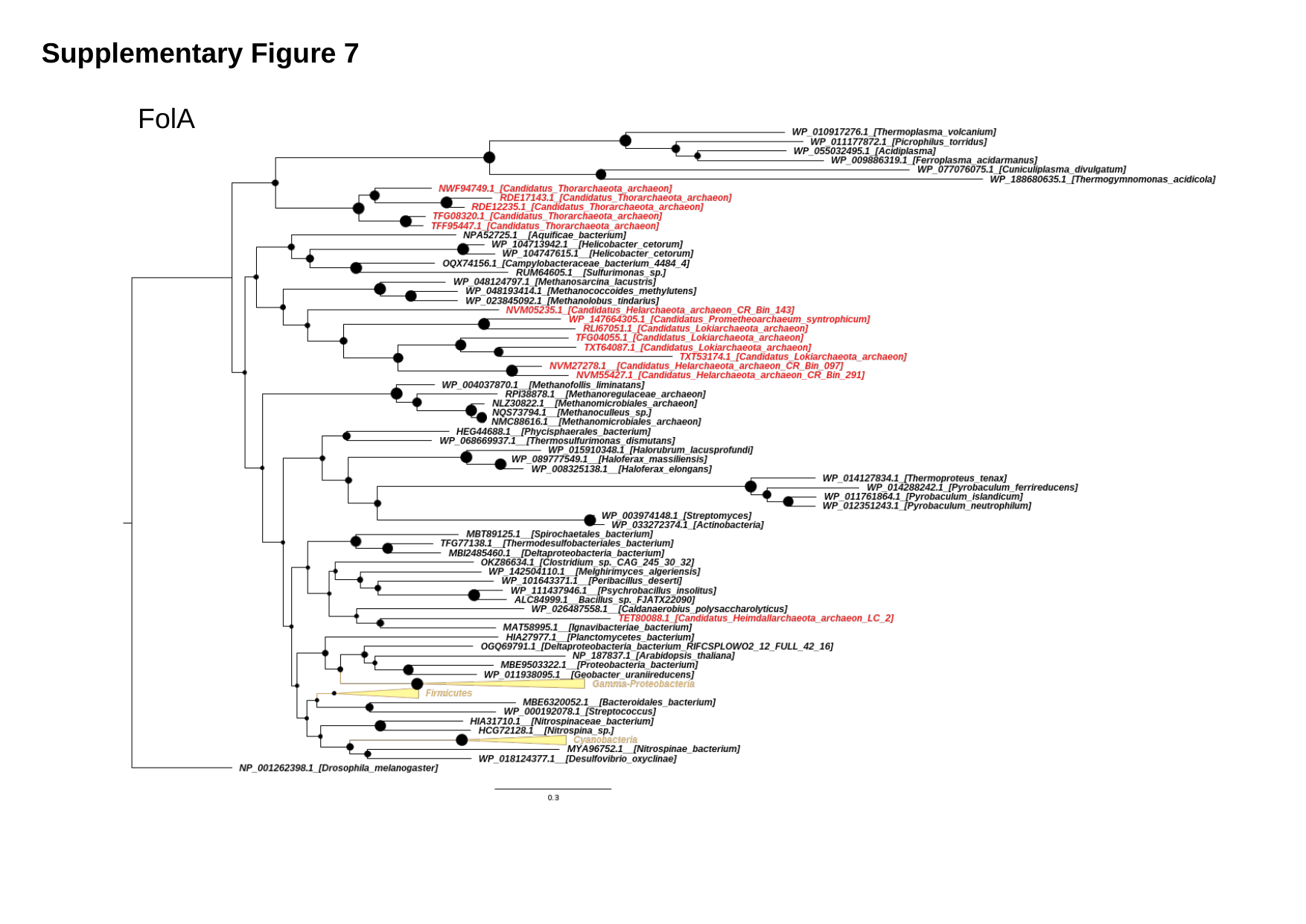

Supplementary Figure 7
FolA

### Slide 9
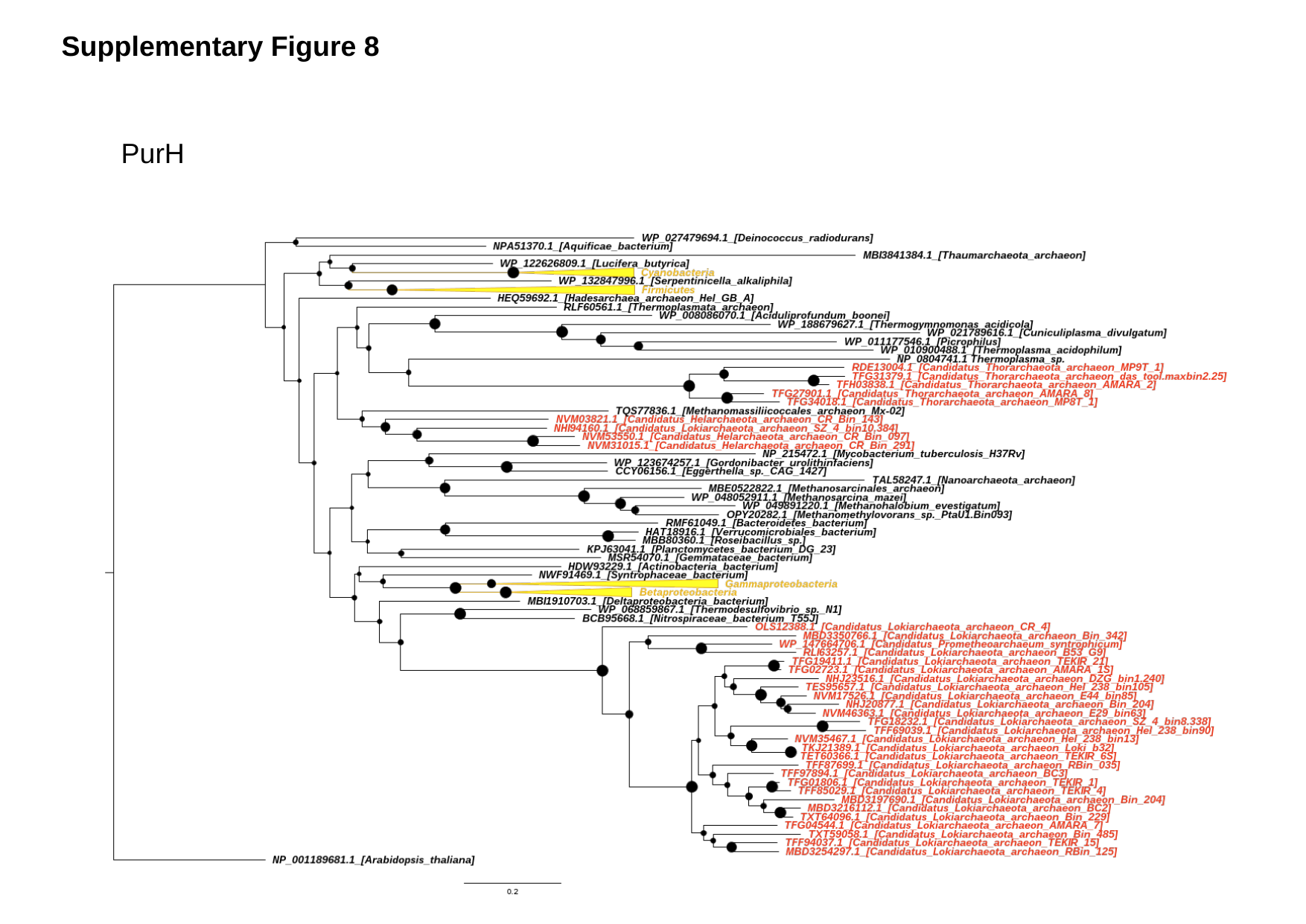

Supplementary Figure 8
PurH

### Slide 10
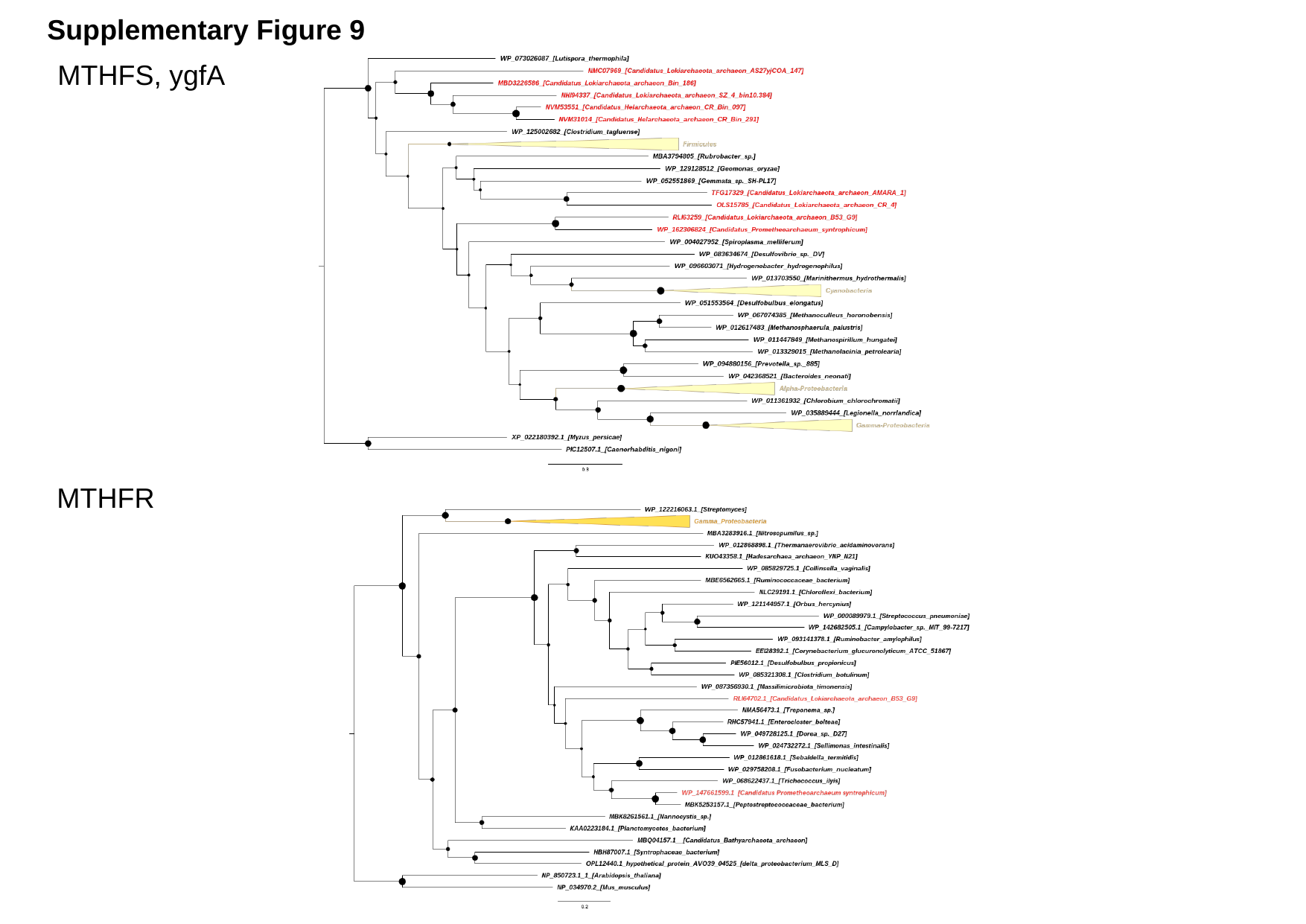

Supplementary Figure 9
MTHFS, ygfA
MTHFR

### Slide 11
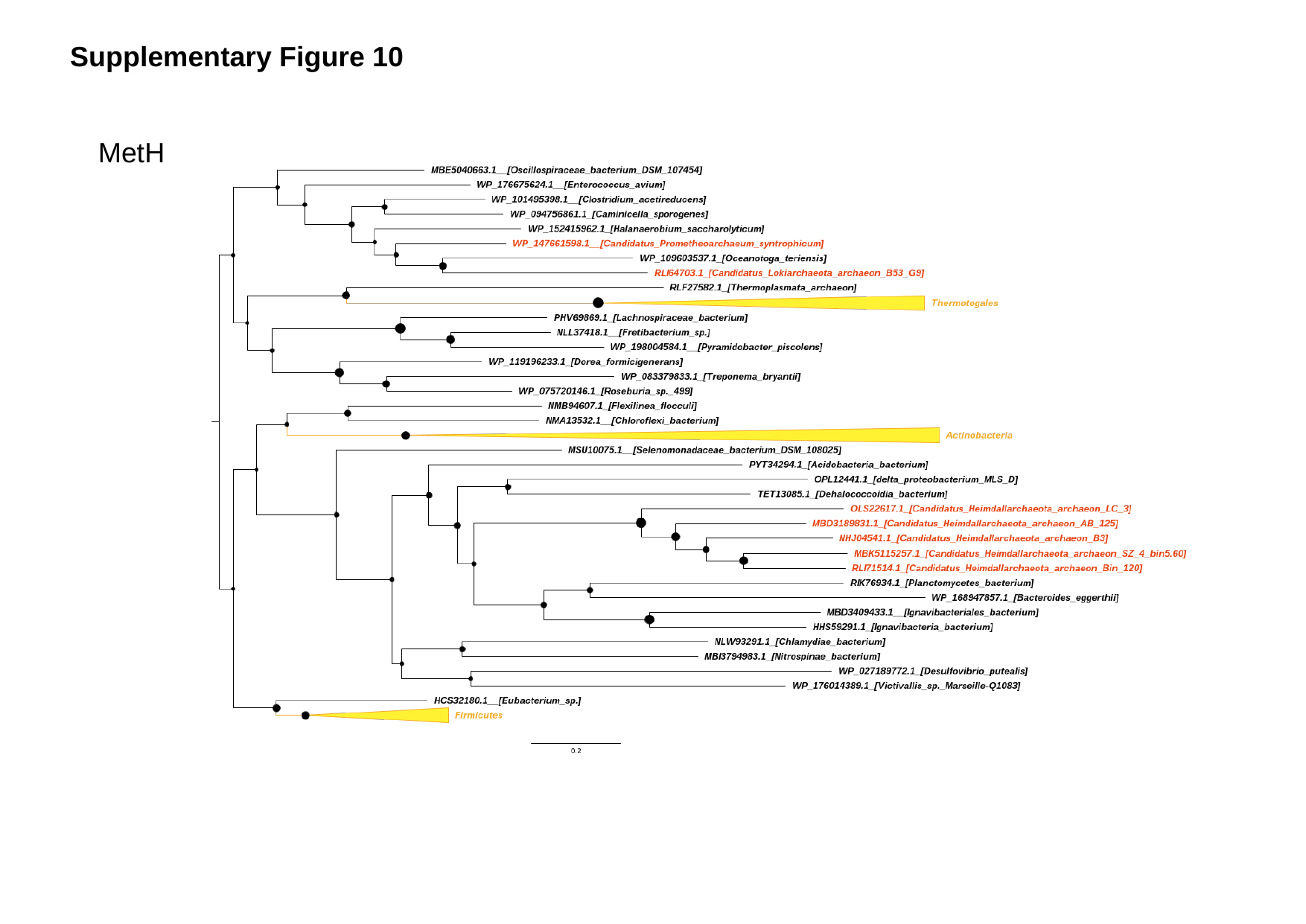

Supplementary Figure 10
MetH

### Slide 12
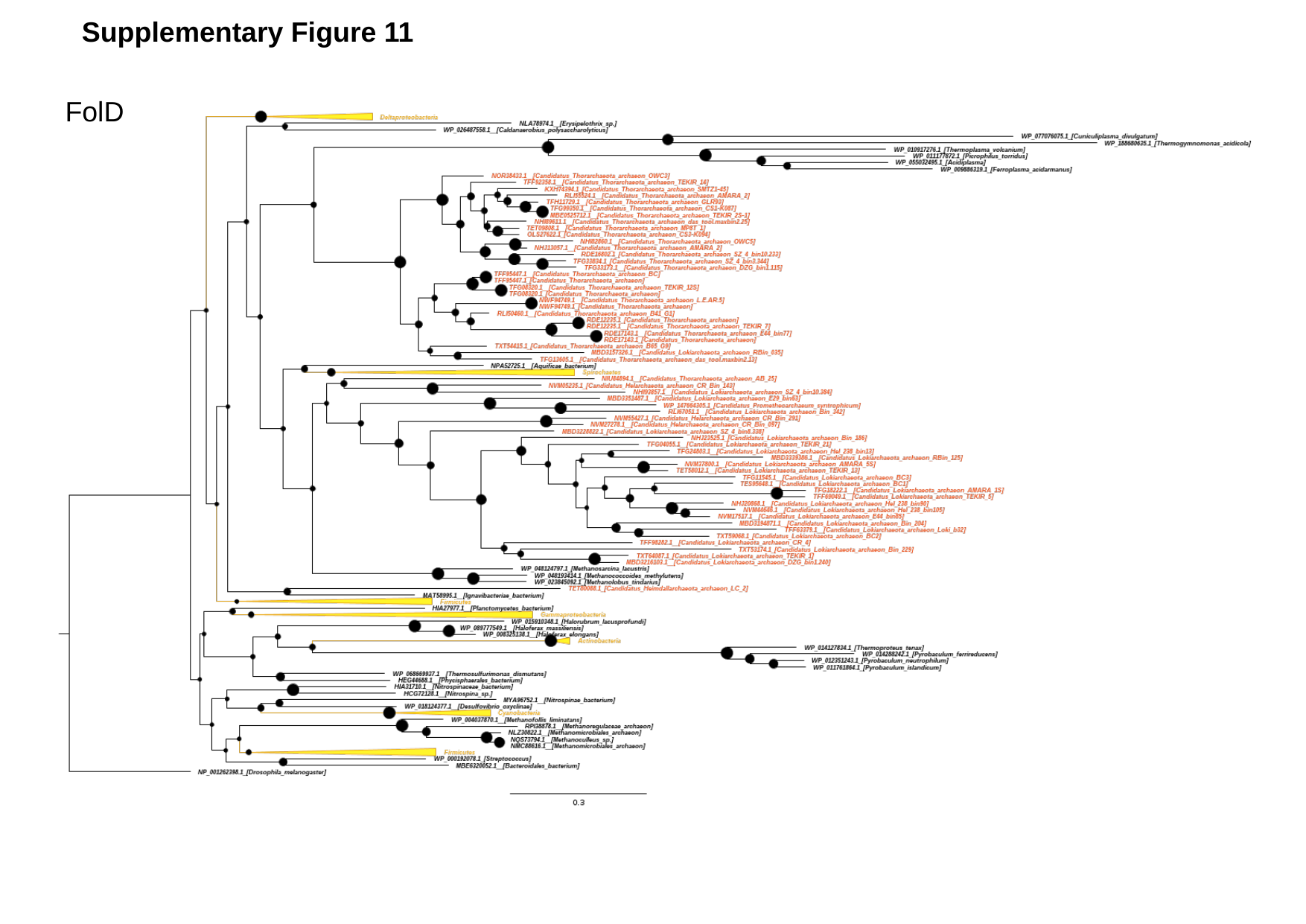

Supplementary Figure 11
FolD
